# A Unified Framework for Self-Recognition Across Monkeys, Infants, and Robots

**DOI:** 10.1101/2025.09.18.677057

**Authors:** Wenqiang Zhang, Kai Chen, Tingyu Zhang, Qianya Sun, Yinqian Sun, Yiyang Jia, Liangtang Chang, Jieqiong Liu, Haoyue Qian, Pengmin Qin, Fan Jiang, Yi Zeng, Guang-Hai Wang, Neng Gong

## Abstract

Self-recognition is a basic cognitive ability that indicates self-consciousness in humans. However, whether non-human agents—such as monkeys and robots—can possess a similar sense of self remains debated. Here, we develop a multi-scenario apparatus to examine self-recognition from different perspectives across macaque monkeys, human infants, and simulated robots. Monkeys trained for mirror-perspective (MP) self-face recognition were able to spontaneously recognize themselves when viewing video images of their back from a third-person perspective (3PP); however, this generalization required a few minutes of “exploration”. In contrast, monkeys trained for 3PP self-back recognition spontaneously demonstrated MP self-recognition within seconds. Meanwhile, first-person-perspective (1PP) self-hand recognition represented a more basic ability that naive monkeys could achieve naturally. Interestingly, human infants aged 1 to 2 years exhibited a stepwise developmental sequence of 1PP, MP, and 3PP self-recognition abilities, consistent with the hierarchical levels of self-recognition in monkeys. Furthermore, we constructed world-model-based simulated robots and found that robots equipped with a body schema exhibited the emergence of self-recognition ability following multisensory integration training, along with the spontaneous generalization of self-recognition across different perspectives. Notably, the formation of internal self-body maps was identified as a key factor enabling this generalization. The convergence of findings across humans, monkeys, and robots not only redefines the evolutionary continuity of the self but also provides an operational framework for designing artificial systems capable of locating themselves in the world.

## Main

The emergence of the sense of self in humans has long been an important subject in cognitive studies. However, the evolutionary origin and neural underpinnings of the self in animals and humans remain unclear. In adult humans, the rubber hand illusion^1^ and out-of-body experience paradigms^2,3^ have been well established for examining body ownership and bodily self-consciousness (BSC)^4^. For infants, the capacity for mirror self-recognition (MSR) typically develops spontaneously around 18 months of age^5^, indicating the emergence of a certain level of BSC. However, the inability of human infants to offer reliable subjective reports during rubber hand illusion and out-of-body experience limits the in-depth study of BSC in infants. Therefore, developing BSC assessment paradigms that are independent of subjective reporting and applicable to infants is especially important for studying the development of self-consciousness in humans. Bodily self-consciousness is also likely to exist in many species besides humans, and experimental paradigms are being developed^6–9^. The MSR using mark test stands as the most extensively employed method for detecting BSC across species. To date, only humans and a small number of great apes are generally considered to be capable of spontaneously passing this test^10–13^. Yet, accumulating evidence suggests that various non-primate animals, including elephants^14^, dolphins^15^, birds^16–18^, even rodents^19^ and fish^20–22^, can also display mirror-induced self-directed behaviors. Macaque monkeys have long been regarded to be incapable of passing the MSR test^23,24^. However, recent studies have shown that macaque monkeys could exhibit MSR behaviors strikingly similar to those of humans after training for association of visual and somatosensory/proprioceptive information^25,26^. These trained monkeys showed typical action of touching marks on their faces and bodies in front of a mirror, together with other mirror-induced self-directed behaviors. However, whether trained monkeys that pass the MSR test truly possess the concept of bodily self remains debatable^27–30^.

Although whether and how artificial intelligence (AI) could possess a sense of self remains controversial, many robotics studies have already addressed the topic of the self, particularly in embodied agents^31^. Developing advanced robots and understanding the human self turn out to pose similar challenges. Leaving aside the development of robots with subjectivity and the associated ethical concerns, advances in robotics can indeed help us understand the scientific nature of the sense of self and, building on that understanding, develop intelligent agents with higher-level cognition. Recently, researchers have employed various robots to investigate diverse self-related behavioral phenomena^31,32,33^. In robotic systems, certain aspects of the self can be systematically constructed by integrating hypothesized components—such as specific brain processes—into embodied intelligent agents and testing them in real-world scenarios. This approach makes it possible to determine whether such components are sufficient and necessary for producing self-related behaviors, thereby offering clues for understanding the self in humans. For example, the sense of body ownership and agency, as fundamental aspects of the self, has been extensively studied in robots, contributing to our understanding of how the brain represents body schema^34,35^.

Paradigms for testing BSC, such as the rubber hand illusion and MSR, have also been investigated in robotic studies through brain-inspired modelling^33,36^. However, the human self is better characterized as a generalizable higher-order cognitive capacity rather than as a simple computational model tailored to specific self-related tasks. To truly understand the human self, it is necessary to develop an operationalizable experimental paradigm for self-generalization that is applicable across biological organisms and intelligent agents.

In this study, we developed a multi-scenario apparatus to examine self-recognition from different perspectives across macaque monkeys, human infants, and simulated robots. First, we found that monkeys could generalize self-recognition across tasks involving different perspectives, including the first-person perspective (1PP), mirror perspective (MP), and third-person perspective (3PP). Furthermore, these self-recognition abilities in monkeys were organized hierarchically across perspectives. Second, in human infants aged 1 to 2 years, we observed a stepwise emergence of 1PP, MP, and 3PP self-recognition abilities, which was consistent with the hierarchical pattern observed in monkeys. Finally, we constructed world-model-based simulated robots and found that robots equipped with a body schema showed emergent self-recognition ability after multisensory integration training, as well as spontaneous generalization of self-recognition across different perspectives. Notably, the formation of internal body maps was identified as a key factor underlying this generalization. This study provides a cross-species and cross-agent framework for understanding the emergence and generalization of self-recognition.

### Multi-scenario self-recognition tasks

To determine whether rhesus monkeys have developed the concept of bodily self, we applied the strategy of examining their ability to demonstrate reasoning and generalization in novel self-recognition tasks besides MSR. Inspired by the rubber hand illusion and out-of-body experience paradigms in human BSC research^1,3^, we designed a self-recognition task apparatus for rhesus monkeys that could be used under various scenarios (Fig. 1a). In this setup, the rhesus monkey sits on a specially designed monkey chair, capable of viewing real-time video images of various body parts, such as its hands, face, and back, on the video display screen positioned in front of the chair. Based on the location of the video camera, we categorized the self-recognition tasks for three scenarios: the first-person perspective (1PP), third-person perspective (3PP), and mirror perspective (MP). In the 1PP self-hand recognition task, the monkey’s hands were placed under an opaque board, and it could only observe the hand movements on the screen, analogous to the paradigm of human rubber hand illusion. In the 3PP self-back recognition task, the monkey could see a video image of its back on the screen, somewhat like the human out-of-body experience paradigm^3^. In the MP self-face recognition task, the monkey could view a mirrored video image of its face on the screen, as in the standard MSR test. Given that rhesus monkeys are unable to provide subjective reports as humans do, we used the mark-touching as a means for the monkeys to indicate self-recognition. The marks used in this study included light spots and dye marks, as well as virtual spiders on the screen, which were used to exclude potential involvement of somatosensation and reflect more instinctive behaviors (Fig. 1a).

**Fig. 1:**
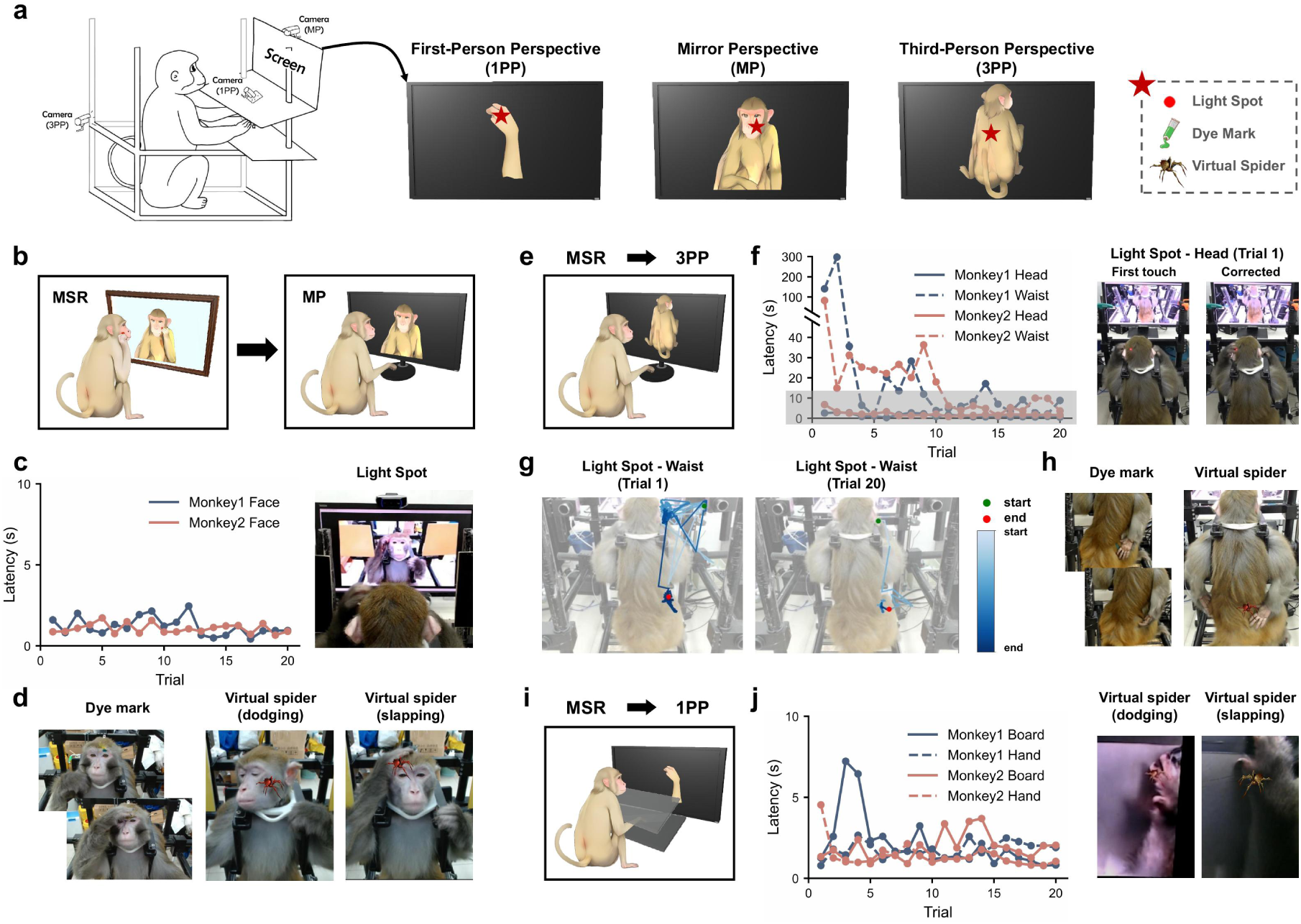
Spontaneous generalization of self-recognition in macaque monkeys. **a,** Apparatus for first-person (1PP), mirror (MP) and third-person (3PP) views, with light-spot, dye-mark and virtual-spider tests. **b,** Transfer from mirror self-recognition (MSR) to the video-based MP test. **c,** Face-touch latencies over the first 20 MP trials in two MSR-trained monkeys, with a representative response. **d,** Representative dye-mark responses and dodging or slapping at virtual spiders in MP. **e,** Switch to a view of the back in 3PP. **f,** Head- and waist-touch latencies over the first 20 3PP trials in the same monkeys. Images show the initial forehead touch and its correction; shading marks the range of rapid responses. **g,** Hand trajectories of Monkey 1 in the first and twentieth waist-target trials. Green and red dots mark the start and end; blue shading indicates movement progression. **h,** Representative 3PP responses to a dye mark and virtual spider on the back. **i,** Switch to the 1PP hand and board view. **j,** Hand- and board-touch latencies over the first 20 1PP trials in the two monkeys, with examples of dodging and slapping responses.

Two rhesus monkeys underwent visual-proprioceptive sensory integration training and demonstrated the ability in passing the standard MSR test (Supplementary Video 1), as shown in the previous studies^26^. Then, these monkeys were placed in the multi-scenario apparatus for testing MP self-recognition (Fig. 1b). Given the high similarity between the MP self-recognition and MSR, both rhesus monkeys immediately (< 3 s) touched the red light spot on the face (Fig. 1c) and also successfully passed the tests involving dye marks and virtual spiders (Fig. 1d, Supplementary Video 1). During the virtual spider test, the monkeys exhibited distinct dodging and slapping behaviors towards the virtual spiders, as if the spiders were physically present on their faces (Fig. 1d, Supplementary Video 1).

### Spontaneous generalization of self-recognition in macaque monkeys

To examine whether monkeys could generalize the ability of self-recognition, we investigated the performance of monkeys capable of MP self-recognition in the 3PP and 1PP tasks. In the 3PP self-back recognition task (Fig. 1e), when a light spot was projected to the back side of the monkey’s head for the first time, the monkey initially touched the forehead, but adjusted the touching on the back side of the heads within 10 seconds (Fig. 1f, Supplementary Video 2). When the light spot was projected to the waist area of monkeys’ backs for the first time (trial 1), they were unable to touch the light spot (Fig. 1f,g) and displayed apparently exploratory hand movements for reaching the light spot on their backs (Supplementary Video 2). After several minutes of “exploration”, both monkeys were able to correctly touch the light spot on their waists by viewing the video image of their back on the screen. After several trials, both monkeys could quickly touch the light spot on their backs (Fig. 1f,g, Supplementary Video 3). Subsequently, they also passed the dye mark and virtual spider tests on their backs (Fig. 1h, Supplementary Video 3). Finally, in the 1PP self-hand recognition task (Fig. 1i), both monkeys promptly passed the test by touching the mark on one hand with the other hand in less than 10 s (Fig. 1j, Supplementary Video 4). These results suggest that monkeys with MSR ability can spontaneously generalize self-recognition to 3PP and 1PP scenarios.

We then inquired whether naive monkeys without MSR training could successfully pass the 3PP and 1PP self-recognition tests. We thus selected two naive monkeys that had only been trained to touch visible light spots in the environment. Given that 1PP self-hand recognition task has been previously reported in rhesus monkeys and is assumed to be relatively straightforward^37^, we first carried out the 1PP self-hand recognition test on these two naive monkeys (Fig. 2a). To assist the monkeys in adapting to the testing apparatus, we placed some food on an invisible board in front of them, and the monkeys could freely reach for the food with their hands (Supplementary Video 5). Simultaneously, the monkeys could observe the real-time video image of their hands grasping the food on the display screen (Fig. 2a, Supplementary Video 5). During the first week, both naive monkeys could not pass the light spot tests on their hands (Fig. 2a, Supplementary Video 5). Nevertheless, the success rate of grabbing food based on the video image gradually increased (Fig. 2a). By the second week, both naive rhesus monkeys began to spontaneously recognize that the hands on the video screen were their own. This was confirmed by their successful touching of the light spot, dye, and spider marks (Fig. 2a, Supplementary Video 5). This finding is in line with previous reports that monkeys can spontaneously identify their own hands after several days^37^. Subsequently, we conducted the 3PP self-back recognition test on these two rhesus monkeys and found that both monkeys failed to pass any mark tests on their backs (Fig. 2b). They did not even exhibit any exploratory behavior towards the light spot that appeared on their backs (Fig. 2b, Supplementary Video 6). Intriguingly, these two monkeys were able to successfully obtain the food placed behind them by viewing the video image (Fig. 2b, Supplementary Video 6), indicating that they could utilize the screen information for spatial mapping, yet they were unable to recognize that the body on the screen belonged to themselves.

**Fig. 2:**
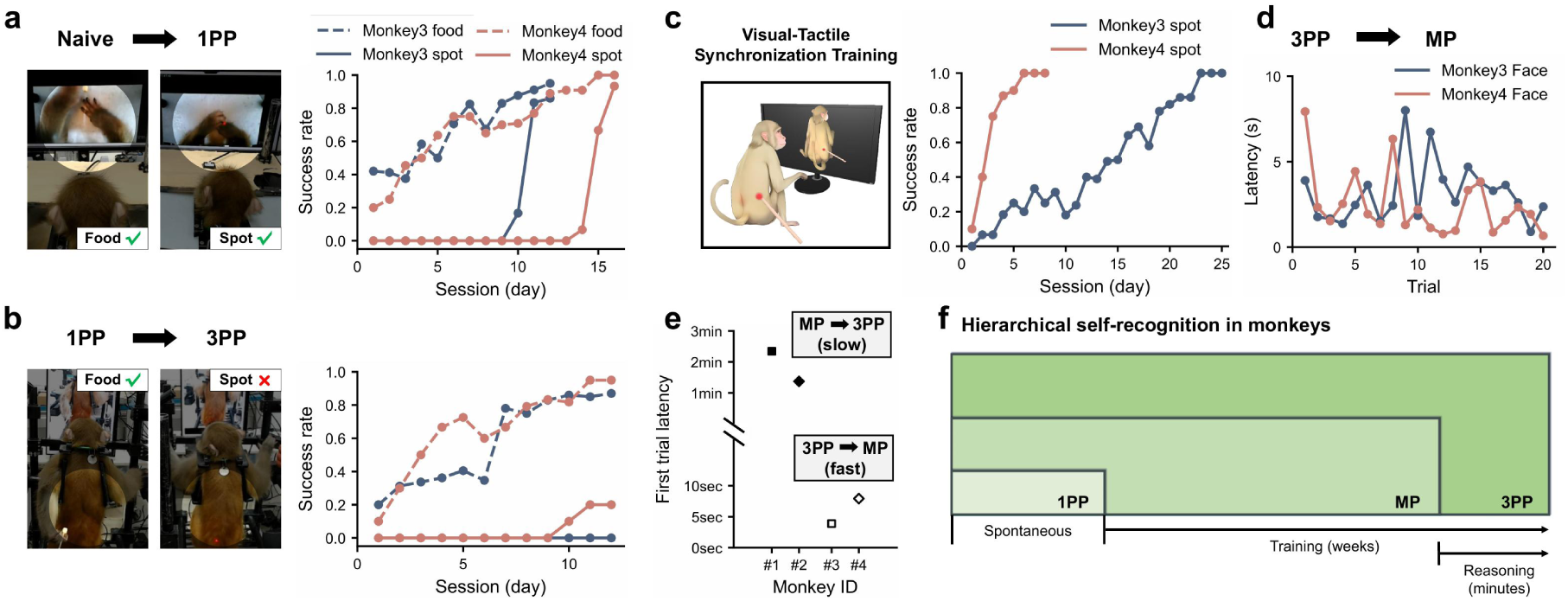
Self-recognition abilities in monkeys showed a hierarchical pattern. **a,** Food retrieval and hand-directed light-spot touching in 1PP, with representative images and success rates by daily session for Monkeys 3 and 4. **b,** Food retrieval and back-directed light-spot touching after switching to 3PP. Curves show success rates by session for the same monkeys. **c,** Visual–tactile training in 3PP. The experimenter touched the back at the light-spot location; curves show spot-touch success by session. **d,** Face-touch latencies over the first 20 MP trials after 3PP training. **e,** First-trial latencies for MP→3PP (Monkeys 1 and 2) and 3PP→MP (Monkeys 3 and 4). Each symbol represents one monkey. **f,** Schematic of the proposed hierarchy from 1PP hand recognition through MP face recognition to 3PP back recognition.

To promote the 3PP self-back recognition in these monkeys, we adopted the visual-tactile synchronization training method used in studying human out-of-body experience^3,4^. Whenever a light spot was projected on the monkey’s back, the experimenter simultaneously used a stick to poke the exact position of the light spot on the monkey’s back, creating a synchronized visual and somatosensory experience (Fig. 2c, Supplementary Video 7). We found that both monkeys were able to pass the back mark tests after such training, with the required training duration apparently depending on the age of the monkey – 7 days and 4 weeks for the younger (aged 6 years) and older (aged 12 years), respectively (Fig. 2c, Supplementary Video 7). Thus, the monkeys could achieve 3PP self-recognition, albeit with a much longer period of synchronized visual and somatosensory experience as compared to that for humans to gain out-of-body illusion (a few minutes)^3^. Subsequently, we adjusted the screen image to the mirrored video image of the face, and found that these two rhesus monkeys passed the MP self-recognition task in less than 10 seconds (Fig. 2d, Supplementary Video 8) and showed MSR behaviors in their home cage equipped with a mirror. This is in sharp contrast to the previous finding^25,26^ that naive rhesus monkeys had to undergo several weeks of training to exhibit MSR ability. We further compared the time required for the monkey to demonstrate self-recognition (during the first trial) after switching the video view from MP to 3PP (Fig. 1f) and from 3PP to MP (Fig. 2d). As shown in Fig. 2e, the time for MP to 3PP switching (> 1 min) was much longer than 3PP to MP switching (< 10 s), indicating the asymmetry in the generalization of self-recognition between the two viewing scenarios. This reveals a hierarchical pattern of self-recognition abilities in monkeys: 3PP self-back recognition requires greater cognitive demand than MP self-face recognition, whereas 1PP self-hand recognition appears to be a more basic form of cognition, which monkeys can achieve spontaneously without training (Fig. 2f).

### Stepwise development of self-recognition in human infants

The rubber hand illusion and out-of-body experience tests are difficult to implement in human infants, but MSR as indicated by face-mark touching has been successfully applied to infants. In this study, we further applied our multi-scenario self-recognition tasks to examine the infants’ ability to perform 1PP, MP, and 3PP self-recognition tasks (Fig. 3a). Twenty-five infants aged between 12 and 24 months were selected for this study after standard physical and cognitive assessments (Extended Data Table 1). Each test was video-recorded, and two experimenters independently evaluated whether the infants passed each of the three tests in the sequence of 1PP, MP, and 3PP, in a manner similar to that the monkey tests, except that the infants were allowed to move in the test room rather than confined to a chair (Fig. 3a). Our results showed a clear developmental trajectory of infants’ self-recognition abilities. Infants were found to be capable of spontaneously passing the 1PP self-hand recognition task around 15 months of age, the MP self-face recognition task by approximately 18 months of age, and the 3PP self-back recognition task at about 23 months of age (Fig. 3b). Furthermore, we selected four infants for in-depth tracking of their development, and found a sequential development of self-recognition in 1PP, MP to 3PP scenarios (Fig. 3c). These findings revealed the development process of human infants’ self-recognition (Fig. 3d). Remarkably, the developmental sequence of self-recognition in three different perspectives aligns well with the above finding of hierarchical levels for self-recognition in rhesus monkeys.

**Fig. 3:**
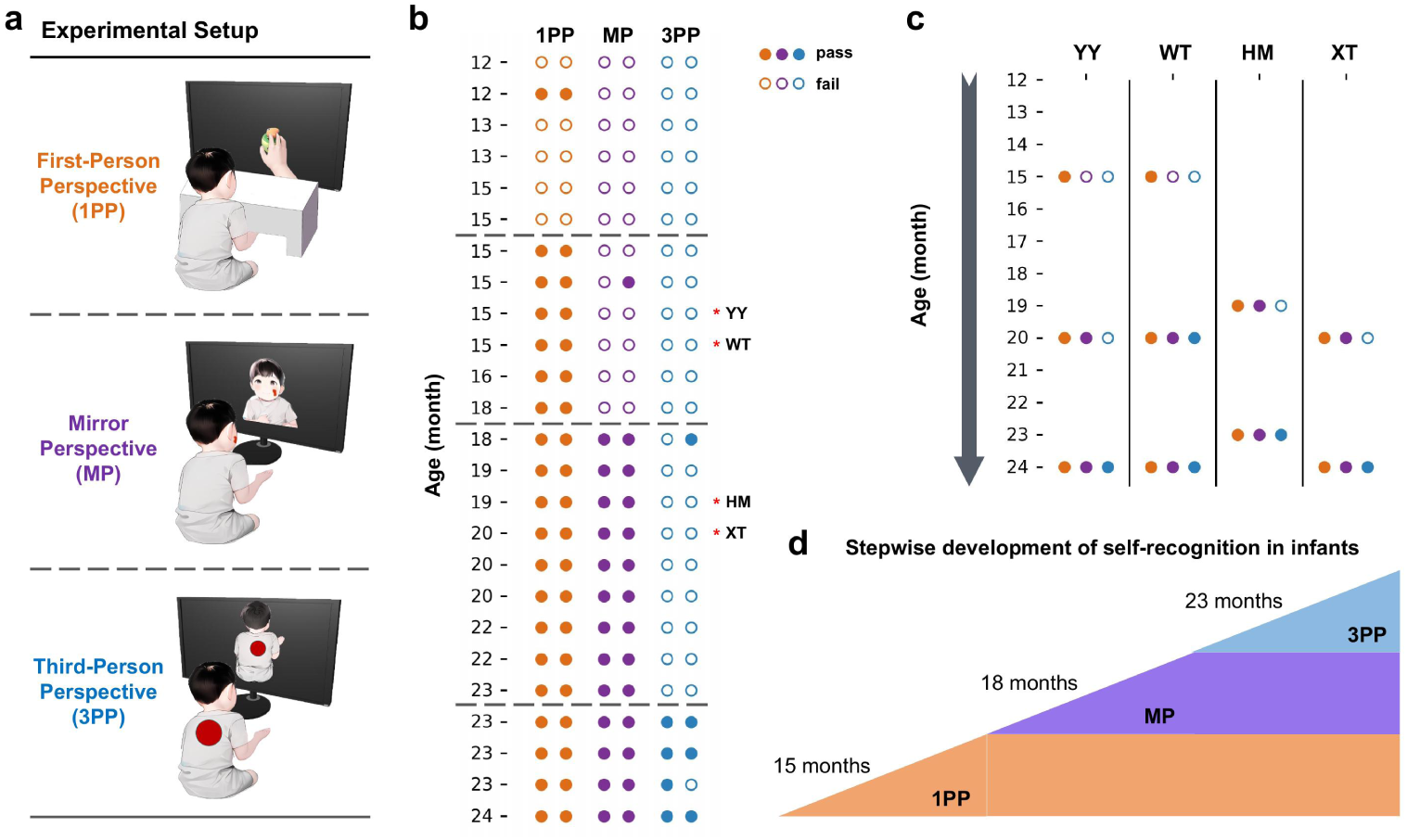
Stepwise development of self-recognition in human infants. **a,** Schematics of the three tasks: using a video of the hand to retrieve a toy in 1PP, touching a face mark in MP and responding to a back sticker in 3PP. **b,** Cross-sectional results for 25 children, ordered by age. Each row is one child; paired circles show independent ratings by two experimenters. Orange, purple and blue denote 1PP, MP and 3PP, respectively. Filled and open circles indicate pass and fail. Dashed lines mark approximate ages of transition; red asterisks identify children followed longitudinally. **c,** Repeated assessments of four children (YY, WT, HM and XT), with one column per child. **d,** Approximate ages of emergence in this sample: 15 months for 1PP, 18 months for MP and 23 months for 3PP.

### Emergence and generalization of self-recognition in simulated robots

We extended our multi-scenario self-recognition tasks beyond biological systems and examined whether and how self-recognition emerges and generalizes in embodied AI systems. We designed MP and 3PP self-recognition tasks for robots in a simulated environment (Fig. 4a). The internal model architecture of the simulated robot was based on the recurrent state space model (RSSM), in which visual, proprioceptive, and tactile information were encoded separately and then integrated into a multimodal module to form latent states used for sensory reconstruction and action control^38^. Building on this basic RSSM model, we added body- and target-position decoders, referred to as the RSSM-body model (Fig. 4b), and then trained the model on a tactile-guided self-touch task without visual input, enabling the model to learn the correspondence among somatosensation, proprioception, and action, thereby constructing a body schema akin to those of humans and monkeys (Fig. 4c). During training, the return of RSSM-body gradually increased and stabilized (Fig. 4d), and the robot was able to quickly and accurately touch target points on its chest or back, while the prediction losses for both body and target positions decreased steadily (Fig. 4e,f). These results indicate that the RSSM-body model can acquire body target localization and the corresponding action control capability without relying on visual input, demonstrating that the robot successfully constructed a body schema.

**Fig. 4:**
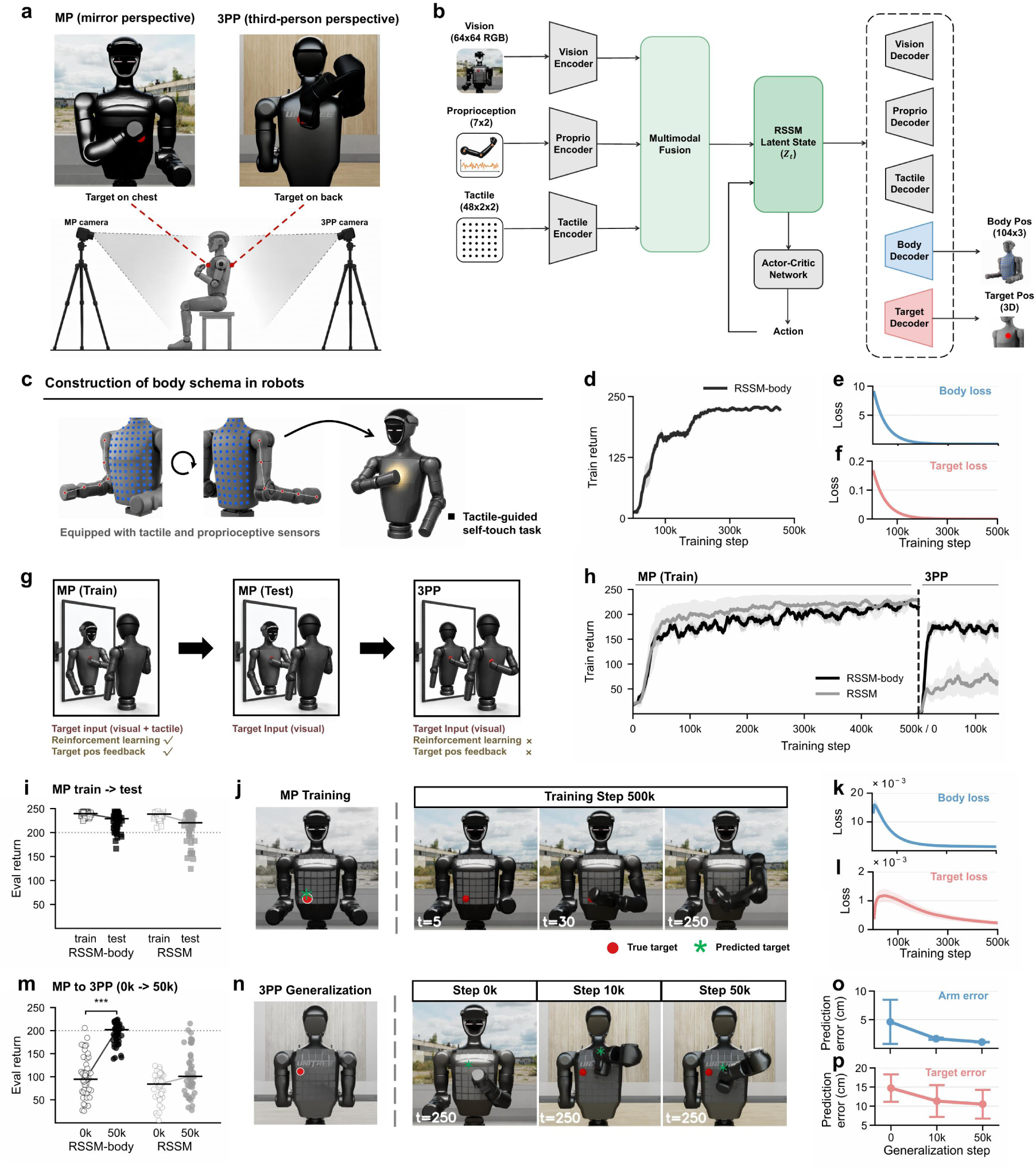
Emergence and generalization of self-recognition in simulated robots. **a,** A simulated Unitree G1 viewed from the MP or 3PP, with example targets on the chest and back. **b,** The recurrent state-space model (RSSM) combines encoded visual, proprioceptive and tactile inputs into a latent state for sensory reconstruction and action control. RSSM-body adds two auxiliary decoders for body landmark and target positions. **c,** Tactile-guided self-touch pretraining without vision. **d,** Pretraining return calculated as the sum of per-frame rewards within each trial (mean and s.e.m. across three seeds). **e,** Body-position prediction loss during pretraining. **f,** Target-position prediction loss during pretraining. **g,** Models first trained in MP with visual targets and tactile stimulation at the target location. Stimulation was removed for MP testing and remained off during 3PP adaptation. During adaptation, reinforcement learning and target-position supervision stopped; only the visual and fusion modules were updated. **h,** Return during MP training and 3PP adaptation (mean and s.e.m. across three seeds). The vertical dashed line marks the switch, where the step count resets. **i,** Eval (Evaluation) returns under MP training and test conditions (open and filled squares), with 50 trials per model and condition. Tactile stimulation at the target location was present during training only. Bars show medians and lines connect them. The dotted line marks a return of 200 in i and m. **j,** Target positions and frames 5, 30 and 250 from a representative RSSM-body trial after 500k MP training steps. Red circles and green asterisks mark true and predicted targets, respectively. **k,** Body-position prediction loss during MP training. **l,** Target-position prediction loss during MP training. **m,** 3PP returns at 0k (open circles) and 50k (filled circles) adaptation steps, with 50 trials per model at each time point. Each point is one trial and bars show medians. Two-sided Mann–Whitney U tests compare returns within each model, with Bonferroni correction for two tests. Stars require adjusted *P* < 0.05 and *** represents adjusted *P* < 0.001. **n,** True target in 3PP (left) and frame 250 of representative trials at 0k, 10k and 50k adaptation steps. Target markers are as in j. **o,p,** Arm (o) and target-position (p) prediction errors at 0k, 10k and 50k adaptation steps (cm; mean ± s.d. across evaluation trials).

We then introduced visual input and examined the emergence and generalization of self-recognition in the robot. First, under the MP scenario, we trained the robot to touch a target that elicited tactile stimulation on its own body in front of a live mirror image, thereby establishing an association between vision and touch, similar to that used in monkeys^25^ (Fig. 4g). During testing, targets that did not elicit tactile stimulation were used, and the robot could only rely on mirrored visual input to touch the target points on its own body (Fig. 4g). Both the RSSM-body and RSSM models were able to learn to localize targets from visual input during training, as reflected by the gradual increase in training return for both models (Fig. 4h). Furthermore, both models exhibited self-touching behaviors and passed the MP self-recognition test with similar evaluation (eval) return (Fig. 4i). As shown in a representative trial at 500k training steps (Fig. 4j), the RSSM-body robot was able to accurately predict the location of the target point after seeing it on its chest and gradually move its hand to the target, with both body- and target-position prediction losses decreasing steadily (Fig. 4k,l). This result also suggests that self-touching of target points from a single perspective can be achieved based on simple computation of spatial position mapping. After completing MP self-recognition training, we switched the visual input to 3PP and stopped reinforcement learning, allowing the model to update only its visual and multisensory integration modules. The model therefore could not accomplish the task by relearning motor strategies, and the tactile stimulation of the target point was removed so that the model had to rely solely on visual information to determine the target location (Fig. 4g). After the perspective switch, the behavioral performance of both models initially declined, exhibiting behavior consistent with touching the chest according to the MP rule even though the target point was actually on the back—similar to the behavior of monkeys after switching from MP to 3PP (Fig. 1f,g).

As generalization proceeded, the RSSM-body model rapidly recovered and maintained a high return, whereas the basic RSSM model remained at a low level (Fig. 4h), and the eval return only in the RSSM-body model increased markedly during generalization (Fig. 4m). As shown in the representative trials (Fig. 4n), the RSSM-body robot gradually shifted from touching the chest to touching the target point on the back, and accordingly, the arm- and target-position prediction errors decreased with increasing generalization steps (Fig. 4o, p). Meanwhile, the results of 3PP self-recognition training and MP self-recognition generalization were exactly the same: both models could complete the 3PP self-recognition task through training, but only the RSSM-body robot could achieve spontaneous generalization in the MP self-recognition task (Extended Data Fig. 1). During generalization, reinforcement learning updates were stopped and target position feedback was removed, thus the RSSM-body model did not relearn target-position prediction and motor control but rather achieved rapid adaptation based on what it had learned before the perspective switch. Therefore, the robot based on the RSSM-body model recapitulated the emergence and generalization of self-recognition observed in monkeys.

### Formation of internal body maps during self-recognition in robots

To investigate the intrinsic mechanism underlying the spontaneous generalization of the RSSM-body model, we compared the latent space representations of the two models during training and generalization. We trained linear decoders to predict the 3D coordinates of body key points and target points from the latent space. During MP self-recognition training, the arm- and target-position decoding errors of the RSSM-body model were close to those of the basic RSSM model (Extended Data Fig. 5), consistent with the finding that the two models performed equivalently during MP self-recognition training. We then analyzed the body topology structure in the latent space representations. Targets that elicited tactile stimulation were presented at 96 locations on the robot’s chest and back, and the latent-space representations of the target points perceived visually and tactilely by the robot were visualized via dimensionality reduction. We found that in the RSSM-body model, the 96 point positions formed a clear spatial structure resembling the chest–back layout of the body; notably, the chest-side structure, which is visible under the MP scenario, was spatially more expanded than the back side, with a body-surface topology correlation coefficient of ρ = 0.77 (Fig. 5a). In contrast, in the latent space of the basic RSSM model, these points were entangled, with ρ = 0.09 (Fig. 5a). We also performed dimensionality reduction on the latent space representations of body posture and target positions during the robot’s self-touching action, and found that during MP self-touching, the RSSM-body model exhibited a mesh-like topology, whereas the basic RSSM model exhibited a more line-like structure (Extended Data Fig. 6a,b). The correlation coefficients between latent space distance and posture distance were 0.77 and 0.95 for the RSSM-body and basic RSSM models, respectively, while the correlation coefficients between latent space distance and target position distance were 0.70 and 0.02, respectively (Extended Data Fig. 6a,b). This indicates that the RSSM-body model coordinated the representations of posture and target position well in the latent space, whereas the basic RSSM model, although able to form a body posture representation, did not form a spatial structure for the target position. The above analyses indicate that during MP self-recognition training, an organizational structure corresponding to the real body-surface spatial relationships—an internal body map—emerged in the latent space of the RSSM-body model.

**Fig. 5:**
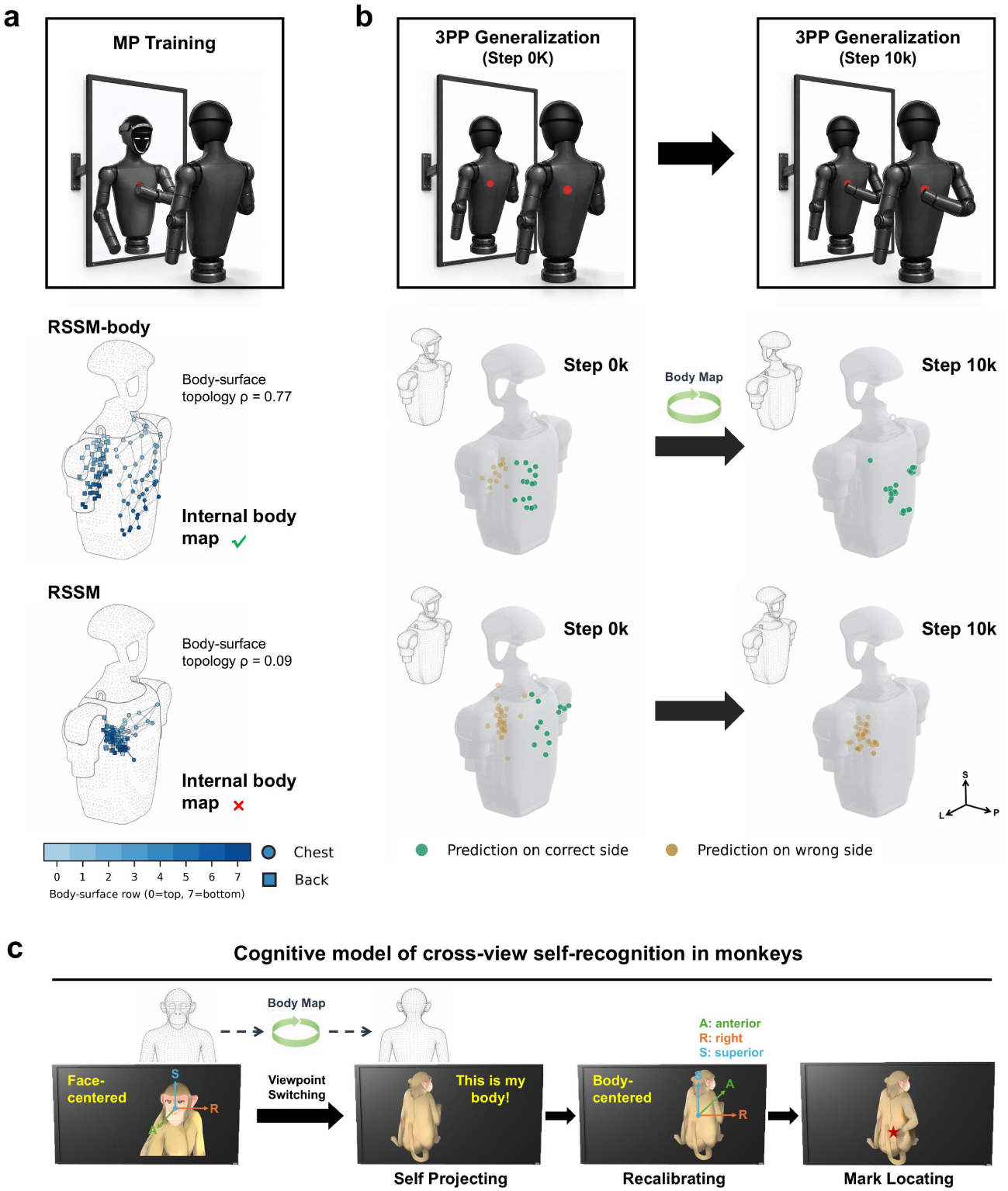
Formation of internal body maps was critical for self-recognition generalization. **a,** Body-surface maps after MP training. With posture fixed, the models received visual input and tactile stimulation at each of 96 torso locations. The points form a three-dimensional embedding of latent cosine distances from multidimensional scaling (MDS), aligned to the torso by Procrustes analysis. Circles and squares mark chest and back locations; blue shades indicate grid rows. Spearman’s ρ is the correlation between latent and physical body-surface distances. **b,** Decoded target positions at 0k and 10k adaptation steps after switching to 3PP. The linear ridge decoder was fitted in MP and held fixed. Each point summarises one trial; green and gold mark predictions on the correct and incorrect body sides, respectively. S, L and P denote superior, left and posterior. **c,** Cognitive model of cross-view self-recognition in monkeys: the animal relates the displayed image to its body, shifts from a face-centred to a body-centred reference frame and recalibrates the mark location. A, R and S denote anterior, right and superior.

We then applied the same linear decoders to the 3PP generalization trials and found that, during the perspective switch, the target-position predictions of both the RSSM and RSSM-body models still predominantly fell on the chest side corresponding to the original MP training (Fig. 5b), consistent with the robot’s behavioral performance. By 10k steps of 3PP generalization, the predictions of the RSSM-body model gradually shifted to the correct back side (Fig. 5b), whereas the basic RSSM still exhibited pronounced chest–back confusion (Fig. 5b). The output coordinate errors of the linear decoders showed that, between 0k and 150k steps of 3PP generalization, the mean arm decoding error of RSSM-body decreased from 6.35 to 4.08 cm, and the target-position decoding error decreased from 12.38 to 8.23 cm, whereas the corresponding errors of the basic RSSM increased from 7.36 to 11.44 cm and from 16.90 to 18.04 cm, respectively (Extended Data Fig. 5). This indicates that, after the perspective switch, only the RSSM-body model successfully recalibrated the body coordinates under the new perspective, and since the target decoder parameters were fixed after MP training, this recalibration achieved by the robot relied on the internal body map formed during MP training. When we analyzed the latent space representations during 3PP self-recognition training and MP generalization, the results were almost identical (Extended Data Figs. 2 and 5). These results indicate that the formation of the internal body map is critical for the robot to acquire a generalizable bodily self-concept.

## Discussion

Bodily self has been normally demonstrated in adult humans by rubber hand illusion and out-of-body experience studies^1–4^. However, there is still debate over whether monkeys could have the concept of bodily self^27–30^. In the present study, we have shown that rhesus monkeys with the ability of MSR or 3PP self-back recognition were capable of spontaneously completing other self-recognition tasks among the three scenarios. These findings strongly indicate the existence of bodily self in trained monkeys, and they can effectively generalize this concept in new self-related tasks. Furthermore, using these multi-scenario self-recognition tasks, we could in-depth examine the developmental emergence of bodily self in human infants, and found a stepwise developmental sequence of 1PP, MP and 3PP self-recognition abilities in infants aged from 1 to 2 years old, in line with the hierarchical levels of self-recognition in monkeys. In the cognitive development process from bodily self to theory-of-mind and narrative self in human children^39,40^, we found that 3PP self-recognition emerges at the time between the emergence of MSR and theory of mind ability, suggesting that recognizing self from 3PP might be a prerequisite for understanding others (theory of mind) and narrative self. It would be of interest to further examine whether these trained monkeys could acquire the ability to understand the behaviors of other conspecifics, similar to that observed in some great apes^41,42^.

Multisensory integration mechanisms during visual-tactile synchronization have been well studied in rubber hand illusion and out-of-body experience studies in adult humans^4,43,44^. In this study, by employing a similar visual-tactile synchronization training method, rhesus monkeys can acquire the 3PP self-back recognition ability. Previous studies have also reported that visual-somatosensory association training could lead to MSR in monkeys^25^, and self-touch in front of mirror could promote MSR in human infants^45^. The multi-scenario self-recognition task we have established offers a useful experimental paradigm for investigating the formation, reasoning, and generalization of the concept of bodily self, as well as the underlying neural mechanisms in rhesus monkeys and human infants. The similarity in the order of emergence of self-recognition implies that the formation of the bodily self in humans and rhesus monkeys may share similar neural underpinnings.

Building robots with a sense of self and understanding the human self share common challenges, which indicates that robotics research can offer clues for studying self-processing in biological organisms^31,46^. Based on the RSSM world model, we extended multi-scenario self-recognition tasks to simulated robots. In order to construct a body schema for a robot (RSSM-body model), we equipped it with somatosensory sensors and trained it with a tactile-guided self-touch task, after which it might have acquired a certain degree of body ownership. Robots with the RSSM-body model could not only learn MP or 3PP self-recognition through multisensory integration training but could also spontaneously generalize self-recognition to new scenarios, just as monkeys did. In contrast, the robot with the basic RSSM model could also learn to touch target points on its own body in a single scenario, suggesting a computation of simple spatial mapping, but it could not achieve generalization. Generalization is a key feature of human self-consciousness and might also serve as a potential criterion for judging whether a robot possesses a bodily self in a genuine sense. During multisensory integration training for self-recognition, we found that the RSSM-body robot formed an internal body map, a factor previously considered important for the sense of body ownership in humans^47^, but whether it is required for modeling the bodily self in robots remains unclear^31^. Our current results demonstrate that the formation of an internal body map is crucial for robots to acquire a genuine bodily self, suggesting a general principle that underlies the bodily self across species and agents.

When switching perspectives, monkeys and robots exhibited similar behavioral manifestations, we thus proposed a cognitive-behavioral model in which monkeys utilize the self-body map for reasoning and generalization (as illustrated in Fig. 5c). First, the monkeys trained in MP or 3PP self-recognition developed the mental self-body map via visual-somatosensory association training. As their body images from different perspectives were presented, they project themselves onto the on-screen body image by rotating the mental self-body map, thereby establishing the self-affirmation of “this is me”, and then proceed to explore the location of the light spot on their own bodies. Given the changes of coordinates between different perspectives, several attempts were required to recalibrate the coordinates during the exploration phase. Once the recalibration is successfully completed, monkeys can precisely locate the mark on their bodies to fulfill the self-recognition task. The apparent variation in the difficulty and the time required for self-recognition among three scenarios suggest relative differences in cognitive demands. Future research into the neural mechanisms underlying self-recognition in monkeys will offer valuable insights into the evolutionary origins of bodily self, as well as provide clues for better constructing and monitoring self-consciousness in embodied artificial intelligence systems.

In summary, we developed a cross-species and cross-agent paradigm to examine self-recognition across different scenarios, and found similar patterns in the emergence and generalization of self-recognition among macaque monkeys, human infants, and simulated robots. These converging findings broaden our understanding of how the sense of self can arise, and provide an empirical framework for studying the bodily self in embodied systems. Moreover, this study offers an operational principle—the formation of an internal body map—for designing artificial systems capable of situating themselves in the world. We wish to clarify that endowing robots with a bodily self-concept is not equivalent to creating artificial systems with subjective experience; nevertheless, the development of robots with more advanced self-models warrants careful attention to ethical issues^48,49^. That said, just as human self-consciousness scaffolds higher-level cognition, an appropriate machine self-concept may scaffold more socially attuned robots. Such systems could better infer others’ intentions and, ultimately, support more harmonious coexistence between humans and artificial intelligence.

## Methods

### Ethics declarations

All animal procedures were approved by the Animal Care Committee of the Shanghai Institutes for Biological Sciences, Chinese Academy of Sciences. The study involving children was approved by the ethics committee of Shanghai Children’s Medical Center (SCMCIRB-K2022187-2). Written informed consent was obtained from a parent or legal guardian before participation.

### Macaques and experimental design

Four healthy male rhesus macaques (Macaca mulatta), aged 6–12 years, were obtained from and housed individually at the Non-human Primate Facility of the Institute of Neuroscience, Chinese Academy of Sciences. Animals were randomly allocated to the two experiments. The two monkeys in Experiment 1 received mirror self-recognition (MSR) training before testing, whereas the two in Experiment 2 had no prior MSR training. The animals received a standard diet, environmental enrichment and water ad libitum. Veterinary staff monitored their health daily.

Before testing, all four monkeys underwent 7 days of habituation and training. They learned to sit in the chair and touch a red light spot projected onto a board to obtain a food reward. Training comprised 200 trials per day. A trial was scored as correct if the first touch was directed to the target location within 5 s of spot onset. Pretraining was considered complete when success exceeded 90% on the final day.

The two monkeys in Experiment 1 had received visual–proprioceptive association training^26^ and passed the standard mirror mark test before the present experiments. They were tested in the mirror perspective (MP), third-person perspective (3PP) and first-person perspective (1PP), in that order. Experiment 2 tested the two monkeys without prior MSR training in the order 1PP, 3PP and MP. In Experiment 2, food-grabbing training took place during the 1PP and 3PP stages, with daily light-spot tests. Dye-mark and virtual-spider tests were performed before and after food-grabbing training. Visual–tactile synchronization training was then introduced in the 3PP stage, followed by assessment of transfer to MP.

### Apparatus for macaque experiments

Each macaque sat in a custom primate chair with its hands resting on a horizontal steel plate (50 × 25 cm). An opaque acrylic panel of the same size was mounted 20 cm above the plate to block the animal’s direct view of its hands. A 27-inch monitor approximately 60 cm in front of the animal displayed real-time video from one of three Logitech cameras operating at 30 Hz.

For the first-person perspective (1PP), a camera mounted centrally on the acrylic panel pointed down toward the hands and the workspace on the plate. For the mirror perspective (MP), a camera above the monitor captured the face and upper body, and the image was horizontally flipped to provide a mirror view. For the third-person perspective (3PP), a camera approximately 60 cm behind the animal captured the head and back. The camera feed was selected according to the test condition.

### Food grabbing and visual–tactile training

Food-grabbing training was conducted only in Experiment 2, under 1PP and 3PP conditions. Food pellets were placed on the steel plate in 1PP and behind the animal in 3PP, where they were visible on the monitor but outside the animal’s direct view. Each daily session comprised 100 trials, with a maximum of 10 s allowed to retrieve the food on each trial. The performance criterion was success above 80% for at least 3 consecutive days.

The two monkeys without prior MSR training received visual–tactile synchronization training during the 3PP stage. A red laser spot was projected onto the head or back while an experimenter gently touched the same location with a stick. Visual and tactile stimulation were presented together for up to 5 s per trial. The macaque received a food reward when it touched the corresponding location on its own body. One session of 100 trials was conducted each day. Training lasted 7 days for the 6-year-old macaque and 4 weeks for the 12-year-old macaque.

### Mark tests across visual perspectives

Before mark testing, each macaque was allowed to observe the real-time video for 5 min. In the light-spot test, an experimenter used a handheld laser pointer to project a red spot at a randomly selected location within the active camera’s field of view. Locations were on the hands or steel plate in 1PP, the head or back in 3PP, and the face in MP. A spot remained visible for up to 3 min, and each session included at least 10 trials. Touching the marked location was scored as a success and rewarded with food to maintain responding across tests.

For the dye-mark test, the monitor was switched off and an odorless, non-toxic green or yellow dye was applied to a randomly selected body location. Plain water was applied at a different location as a control. The animal was observed for 10 min with the monitor off to check for touching caused by applying the dye or water. The monitor was then switched on and behavior was recorded for a further 10 min. Each test was performed at least twice on each testing occasion.

For the virtual-spider test, a custom script using opencv-python superimposed an animated spider on a selected body location in the live video. After several seconds, the spider moved to another location. Dodging, slapping and other behavioral responses were recorded during a 1-min test. Each test was performed at least twice on each testing occasion.

### Child participants and testing procedures

Twenty-five typically developing children aged 12–24 months participated (13 boys and 12 girls). Families were recruited through online announcements from Shanghai Children’s Medical Center. They received a developmental assessment report and either a children’s picture book or 50 yuan as compensation. Children were tested with a parent present in a child-friendly room. Each testing session lasted 1 h and included the 1PP, MP and 3PP tasks in that order.

At the 1PP station, an experimenter and the parent used speech and gestures to encourage the child to put their hands into the box and observe the resulting movements on the monitor. After several minutes of familiarization, an experimenter placed a toy inside the box and recorded whether the child used the video to retrieve it.

At the MP station, the child watched their image for several minutes. The monitor was then switched off and a red dye mark was applied to the face. After checking for spontaneous touching, the experimenter switched the monitor on and recorded the child’s response to the visible mark.

At the 3PP station, the child was encouraged to play while watching the video of their body from behind. After familiarization, an experimenter placed a circular red sticker, approximately 15 cm in diameter, on the parent’s back. The parent guided the child to find the sticker both on the monitor and on the parent’s back. Once the child had done so, the experimenter covertly placed an identical sticker on the child’s back. The child was again prompted to look for the sticker, and their response was recorded. Four children underwent repeated assessments. YY and WT were each tested at 15, 20 and 24 months, HM at 19 and 23 months, and XT at 20 and 24 months. The 1PP, MP and 3PP tasks were assessed at each session.

### Apparatus for child experiments

Testing took place at three stations in one room. Each station had a monitor and a Logitech camera operating at 30 Hz. As in the macaque setup, the monitor was positioned approximately 60 cm from the child and displayed real-time video; the MP image was horizontally flipped to provide a mirror view.

At the 1PP station, the child sat on a small stool at a table holding a white polypropylene box (60 × 30 × 22 cm). The child could put their hands through an opening in the lower part of the box to reach toys inside. A camera on top of the box recorded the interior and sent the image to a monitor in front of the child. At the MP station, the child sat on a stool facing a monitor and camera that provided a live, mirror-like image of the face. At the 3PP station, the child could stand, walk or sit on a rocking horse within a designated area. A camera behind the child recorded a view of the back, which was displayed on a monitor in front of the child.

### Developmental assessments

Development was assessed with the Chinese versions of the Griffiths Development Scales (GDS-C, 0–2 years) and the Ages and Stages Questionnaires, third edition (ASQ-3). Participant characteristics and developmental scores are reported in Extended Data Table 1.

Two trained developmental assessors administered the GDS-C. The locomotor subscale (Scale A) was used to assess gross motor development, including balance and coordination. Its developmental quotient, termed AQ, was calculated as follows.

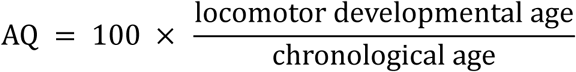

Scores were classified as normal (AQ ≥ 85), mildly impaired (70 ≤ AQ < 85), moderately impaired (55 ≤ AQ < 70) or severely impaired (AQ < 55). All children were classified within the normal range.

The child’s primary caregiver completed the age-appropriate Chinese ASQ-3 questionnaire for ages 12–24 months. ASQ-3 covers communication, problem solving, personal-social skills, gross motor skills and fine motor skills. Each domain contains six items scored as 0 (no), 5 (sometimes) or 10 (yes), giving a domain score of 0–60. All children were classified within the normal range for gross motor development according to the age-specific norms.

### Behavioral analysis

Behavior was scored from video recordings. In the macaque light-spot test, latency was measured manually from the onset of the red spot to a successful touch at the target location. Two experimenters independently rated performance as pass or fail in the children’s tasks. Touch latencies were recorded for individual trials, and success rates were calculated for each testing session. Cross-sectional analyses compared children’s task outcomes across ages; longitudinal analyses compared repeated assessments within each child.

Hand movements during the macaque 3PP light-spot test were tracked from 30-Hz videos using DeepLabCut v2.3.10. Both shoulders, elbows and hands were manually labeled in 17 frames. A ResNet-50 network was trained on 16 frames, with one frame reserved for testing. Predictions used the 27,000-iteration checkpoint after label refinement. Mean landmark localization errors were 2.73 pixels on the training frames and 11.66 pixels on the test frame. Coordinates were smoothed using the DeepLabCut ARIMA filter with order (5, 0, 2). For representative trajectories from trials 1 and 20, right-hand positions with a DeepLabCut likelihood greater than 0.5 were retained and connected in temporal order. Trajectories were plotted in two-dimensional image coordinates, in pixels.

### Simulation and world models

A Unitree G1 humanoid simulated in Isaac Lab v2.3.0 learned to touch targets on its chest or back (Fig. 4a). Each action specified three translational and three rotational increments of the right hand, scaled by 0.02 m and 0.05 rad, respectively. Damped least-squares inverse kinematics converted these commands into movements of seven right-arm joints. Each action command was held for two simulation steps. Each trial contained 250 commands (500 simulation steps). Training and adaptation steps in the figures refer to simulation steps.

Visual input consisted of frontal mirror-perspective (MP) or rear third-person-perspective (3PP) images, resized and padded to 64 × 64 pixels. Proprioceptive input consisted of seven joint angles and their velocities. Angles were divided by π, and velocities were scaled as tanh (0.1v). Tactile input comprised 96 contact-force channels from two 8 × 6 chest and back sensor grids, and 96 channels indicating the sensor receiving the target stimulus. Rectified forces were scaled as 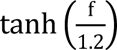. Reward combined hand–target proximity with an action-smoothness penalty. For distance d in meters and action vector a:

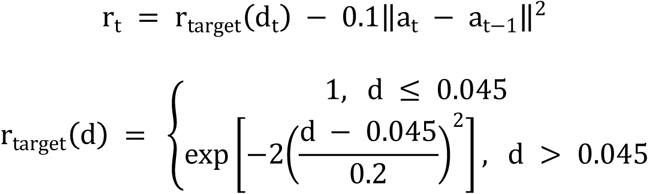

The world model used a Dreamer-style recurrent state-space model (RSSM; Fig. 4b)^38^. A convolutional neural network encoded images, while a multilayer perceptron encoded tactile and proprioceptive inputs. Their outputs were concatenated. A gated recurrent unit updated the recurrent state using the previous latent state and action. The recurrent state was combined with the encoded sensory inputs to infer the current stochastic state, giving its posterior distribution. A separate network predicted the prior distribution before the current observation was incorporated. The 512-dimensional recurrent state h_t_ and 32 categorical variables z_t_, each with 32 categories, together formed 1,536 latent features.

Convolutional and multilayer perceptron decoders reconstructed images and nonvisual signals, respectively. Heads predicted reward and whether the trial would continue. The actor generated actions, and the critic estimated future cumulative reward. Actor–critic learning used 15-step trajectories imagined by the world model, with discount 0.997 and λ = 0.95. RSSM-body added two auxiliary decoders predicting the positions of 104 body landmarks and the target position, while the baseline RSSM omitted them. Landmarks comprised seven arm links, the end effector and 96 torso sensor centers. The visual encoder used convolutional depth 32 and kernel size 4. Nonvisual encoder and decoder networks had five hidden layers of 1,024 units; auxiliary decoders had three of 512 units, and actor and critic networks each had two of 512 units. Networks used SiLU activations and layer normalization.

### Training and cross-view adaptation

Training began with tactile-guided self-touch without vision (Fig. 4c). On each trial, a target stimulus was assigned to the chest or back with equal probability, and the robot learned to move its hand to that location using tactile and proprioceptive input. The image input was set to zero, and the visual modules were frozen.

Visual training added MP or 3PP images while the robot continued the target-touching task. Each target was marked red and elicited tactile stimulation at the corresponding body location. To retain self-touch behavior, 35% of trials placed the target on the side outside the camera view: the back for MP and the chest for 3PP. The nonvisual encoder, nonvisual decoder and body decoder remained frozen during this stage.

Cross-view adaptation switched an MP-trained model to 3PP images, or a 3PP-trained model to MP images (Fig. 4g; Extended Data Fig. 1a). Tactile stimulation at the target location was removed, and actor–critic updates stopped. The visual encoder, visual decoder and observation-conditioned posterior mapping were updated; recurrent transitions, nonvisual pathways and prediction heads remained fixed. Reward, continuation and target-position prediction losses were disabled. Adaptation used image and nonvisual reconstruction, body-position prediction and Kullback–Leibler (KL) losses. Gradients through the fixed body decoder trained the visual encoder and posterior mapping. Rewards were recorded to assess task performance.

The world-model loss combined prediction losses with dynamics and representation KL terms:

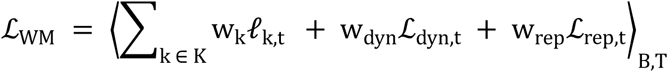

Angle brackets denote averaging over batch and time. Each *P*_k,t_ is a prediction loss for output k at step t. The set K comprises image, tactile, proprioceptive, reward, continuation, body-position and target-position outputs. Images used squared-error loss; vector outputs used squared error after a symmetric logarithmic transformation. Reward and continuation used negative log-likelihood losses. With q_t_ and p_t_ denoting the posterior and prior distributions over z_t_, the KL terms were:

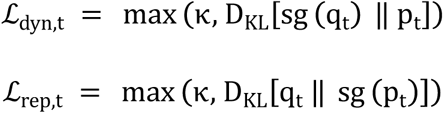

The operator sg stops gradients: the dynamics term trains the prior to match the posterior, and the representation term regularizes the posterior toward the prior. Each KL term had a lower bound of κ = 1. Prediction weights w_k_ and KL weights w_dyn_and w_rep_ were tuned separately for the two models using several sets of values; disabled terms had zero weight.

Training used Adam, world-model and actor–critic learning rates of 10^−4^ and 3 × 10^−5^, batches of 32 sequences of 64 observations, and a replay buffer holding up to 10^6^ transitions. The training ratio was 512 observation steps sampled from replay per newly collected action. Adaptation used a learning rate of 10^−4^.

### Spatial decoding

Auxiliary decoder predictions were compared with simulator coordinates. Arm evaluation used seven arm links and the end effector. After excluding the first five frames, the distance between each predicted and true arm landmark was calculated in every remaining frame. The median of all these distances was taken as the arm prediction error for that trial. Target prediction error was the median distance between predicted and true target positions across the remaining frames. All errors were expressed in centimeters.

To compare spatial information in the latent states of both models, separate linear decoders were fitted using multi-output ridge regression for each model, random seed and training stage. For each decoder, 50 evaluation trials were divided into 35 training and 15 test trials. The first five frames were excluded. Features were standardized and coordinate outputs centered using training data alone; the ridge penalty was α = 1.

Linear decoders were trained on latent states after nonvisual self-touch, MP training or 3PP training and tested on each of these stages. Cross-view testing comprised MP→3PP and 3PP→MP at 0k and 150k adaptation steps, giving three training sources and seven test conditions for each model and for arm and target coordinates. Decoder weights and preprocessing remained fixed during testing. Trial-level errors were first averaged within each random seed and then across seeds.

Spatial plots used the MP-trained target decoder for MP→3PP transfer and the 3PP-trained target decoder for 3PP→MP transfer, comparing 0k with 10k adaptation steps. Each point represented one trial. Its position was calculated by taking the median decoded value for each coordinate across frames. The nearest torso sensor determined whether the predicted target lay on the chest or back.

### Latent spatial geometry

Body-surface representations were examined at 96 sensor-centered target locations while posture was fixed and actions were set to zero. The robot received visual, tactile and proprioceptive signals, including tactile stimulation at the target location on the body. For each target, recurrent state was reset and the same observation repeated for 32 steps. Each of the final 16 latent vectors was normalized; their mean was normalized again to obtain one representation per target. Pairwise cosine distances formed a 96 × 96 matrix, averaged across three seeds.

Classical multidimensional scaling (MDS) embedded the averaged latent distance matrix in three dimensions, with one point per target location. Procrustes alignment translated, rotated or reflected, and uniformly scaled the point cloud to match sensor positions for display. Physical distances were shortest paths on a graph joining adjacent sensors within each grid and linking the chest and back grids at their lateral boundaries. Edge lengths equaled sensor separation. Spearman correlation between the upper triangles of the original latent and physical distance matrices quantified correspondence.

Frame-by-frame analysis used evaluation trials generated by the learned policy. Frames 5, 10, …, 250 from nine target trials yielded 450 observations per model, view and seed. Joint angles were standardized within each view. Spearman correlations compared latent cosine distances with Euclidean distances between standardized postures or target positions. For visualization, classical MDS embedded the latent distance matrix in two dimensions, with one point per frame. Procrustes alignment matched it to a two-dimensional MDS embedding of posture distances. Points were colored by the first principal component of the seven joint angles or by target-grid location. All three seeds were analyzed, and seed 0 was displayed.

### Statistical analysis

Evaluation return was the sum of per-frame rewards within one trial. Two-sided Mann–Whitney U tests compared returns before and after adaptation within each model. Within each transfer direction, P values were Bonferroni-corrected for the two before–after tests, one for each model. Statistical significance required both an adjusted P < 0.05 and an absolute Cliff’s delta of at least 0.474. Analyses used Python, NumPy, SciPy and scikit-learn.

## Data availability

The data supporting this study will be publicly available on Zenodo upon publication. Editors and reviewers can access the data at https://zenodo.org/records/24130593.

## Code availability

The code supporting this study will be publicly available on Zenodo upon publication. Editors and reviewers can access the code at https://zenodo.org/records/24129450.

## Acknowledgments

We thank Mu-ming Poo for helpful discussions and insightful comments on this manuscript.

## Author contributions

N.G., G.W., Y.Z. and F.J. conceived and designed the study. W.Z., K.C., L.C. and H.Q. performed macaque monkey experiments. T.Z., Q.S. and J.L. performed human infant experiments. W.Z., Y.S. and Y.J. performed robot experiments. W.Z., K.C. and T.Z. analyzed data and prepared the figures. P.Q. contributed to the interpretation of the results. N.G. and W.Z. wrote the manuscript. All authors have read and approved the manuscript submission.

## Extended data figures and tables

**Extended Data Fig. 1:**
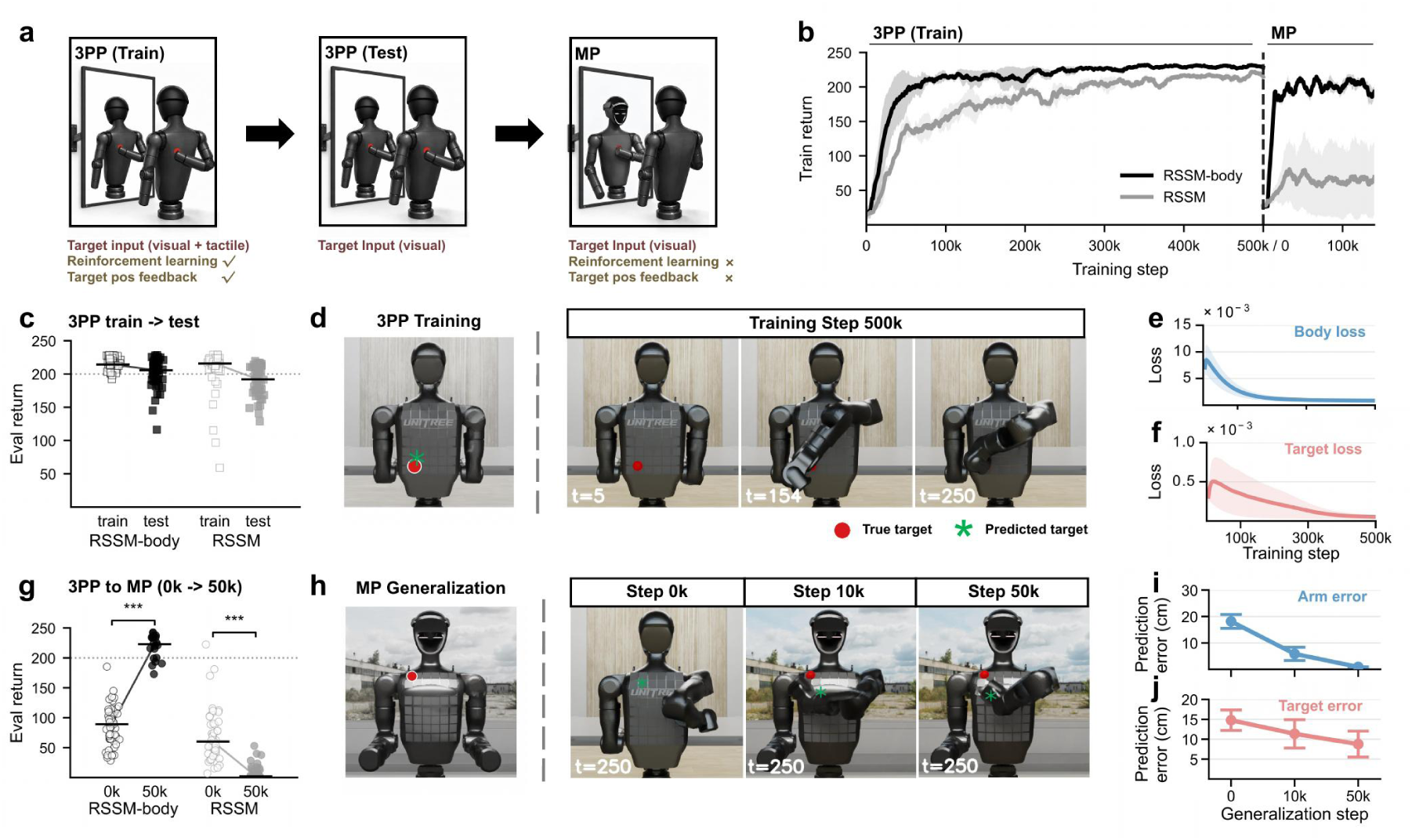
Generalization of self-recognition from 3PP to MP in robots. **a,** Models first trained in the third-person perspective (3PP) with visual targets and tactile stimulation at the target location. Stimulation was removed for 3PP testing and remained off during adaptation to the mirror perspective (MP). During adaptation, reinforcement learning and target-position supervision stopped; only the visual and fusion modules were updated. **b,** Return during 3PP training and MP adaptation (mean and s.e.m. across three seeds). The vertical dashed line marks the perspective switch and step reset. **c,** Eval (Evaluation) returns under 3PP training and test conditions (open and filled squares), with 50 trials per model and condition. Tactile stimulation at the target location was present during training only. Bars show medians and lines connect them. The dotted line marks a return of 200 in c and g. **d,** True and predicted targets and frames 5, 154 and 250 from a representative RSSM-body trial after 500k 3PP training steps. Red circles and green asterisks mark true and predicted targets, respectively. **e,** Body-position prediction loss during 3PP training. **f,** Target-position prediction loss during 3PP training. **g,** MP returns at 0k (open circles) and 50k (filled circles) adaptation steps. RSSM-body had 50 trials at 0k and 20 at 50k; RSSM had 50 at both time points. Points represent trials and bars show medians. Two-sided Mann–Whitney U tests compare returns within each model, with Bonferroni correction for two tests. Stars require adjusted *P* < 0.05 and *** represents adjusted *P* < 0.001. **h,** True target in MP (left) and frame 250 of representative trials at 0k, 10k and 50k adaptation steps. Target markers are as in d. **i,j,** Arm (i) and target-position (j) prediction errors at 0k, 10k and 50k adaptation steps (cm; mean ± s.d. across evaluation trials).

**Extended Data Fig. 2:**
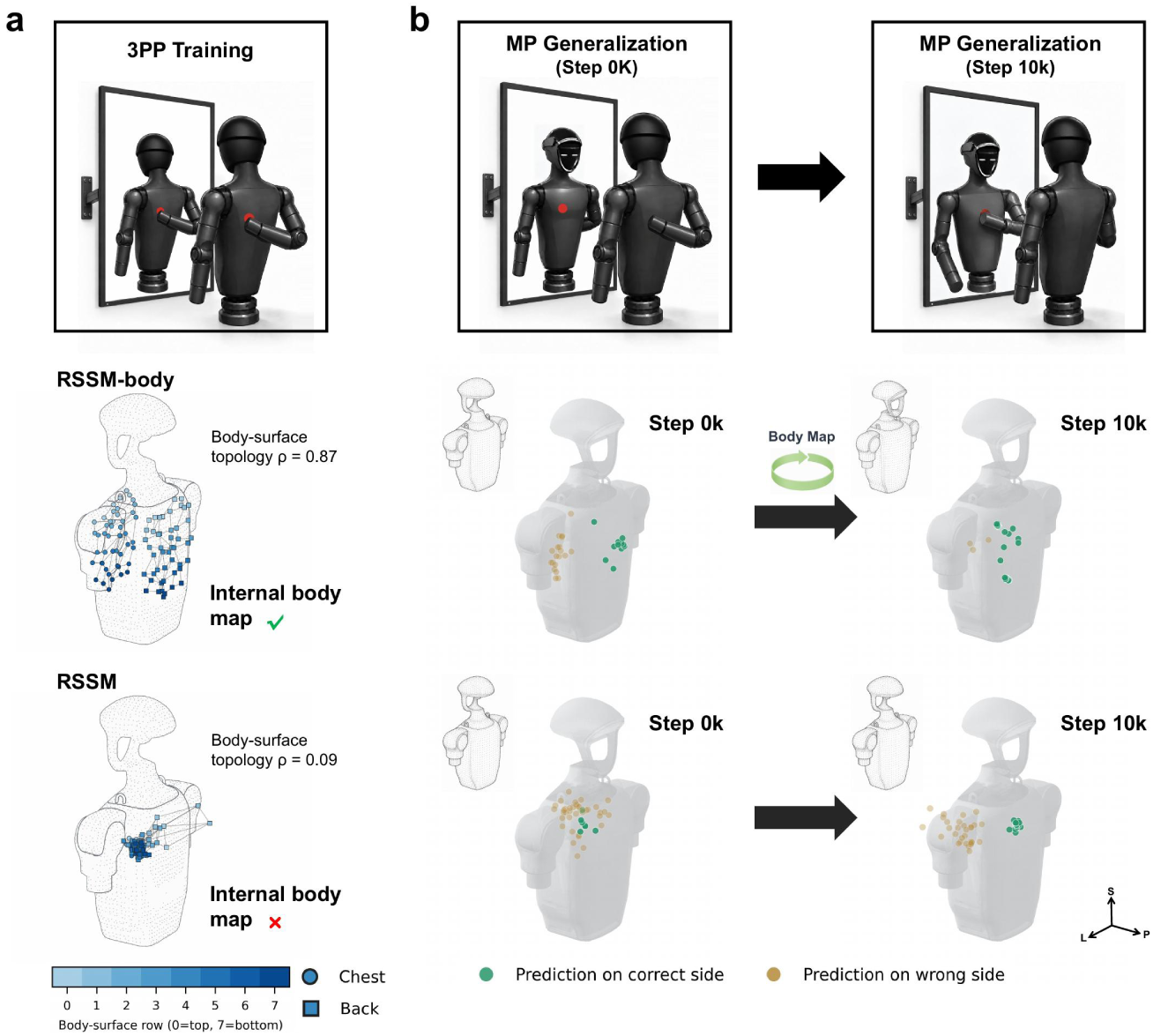
Formation of internal body maps during 3PP-to-MP self-recognition generalization in robots. **a,** Body-surface maps after 3PP training. With posture fixed, the models received visual input and tactile stimulation at each of 96 torso locations. The points form a three-dimensional embedding of latent cosine distances from multidimensional scaling, aligned to the torso by Procrustes analysis. Circles and squares mark chest and back locations; blue shades indicate grid rows. Spearman correlations with physical body-surface distances were 0.87 for RSSM-body and 0.09 for RSSM. **b,** Decoded target positions at 0k and 10k adaptation steps after switching to MP. The linear ridge decoder was fitted in 3PP and held fixed. Each point summarises one trial; green and gold denote predictions on the correct and incorrect body sides. S, L and P denote superior, left and posterior.

**Extended Data Fig. 3:**
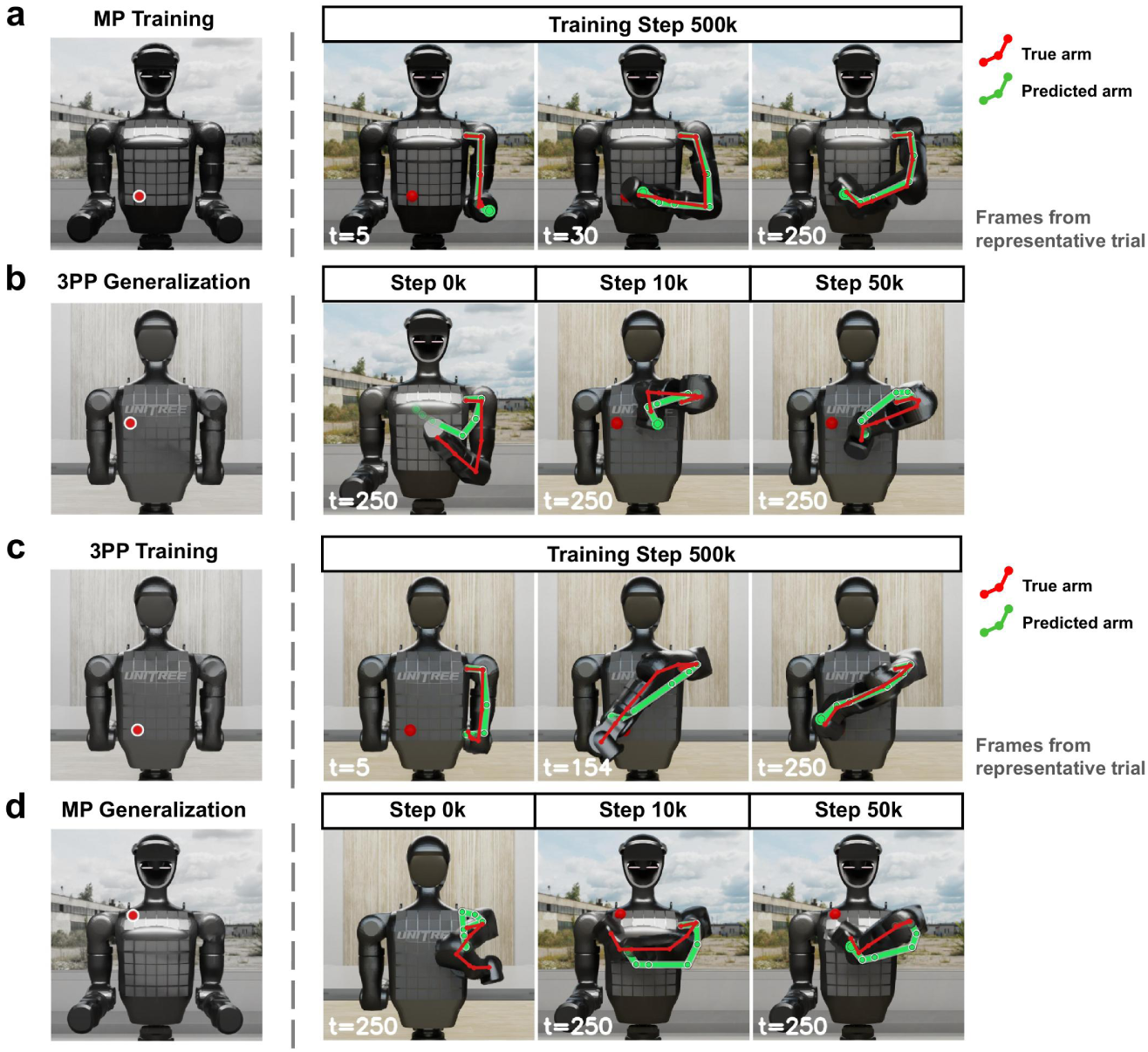
Arm posture prediction during self-recognition training and generalization in robots. **a,** True and predicted arm postures for RSSM-body at frames 5, 30 and 250 of a representative trial after 500k MP training steps. **b,** Frame 250 of representative trials at 0k, 10k and 50k adaptation steps after switching from MP to 3PP. **c,** Frames 5, 154 and 250 of a representative trial after 500k 3PP training steps. **d,** Frame 250 of representative trials at 0k, 10k and 50k adaptation steps after switching from 3PP to MP. The left image in each row shows the target condition; red and green joint chains show true and predicted arm postures.

**Extended Data Fig. 4:**
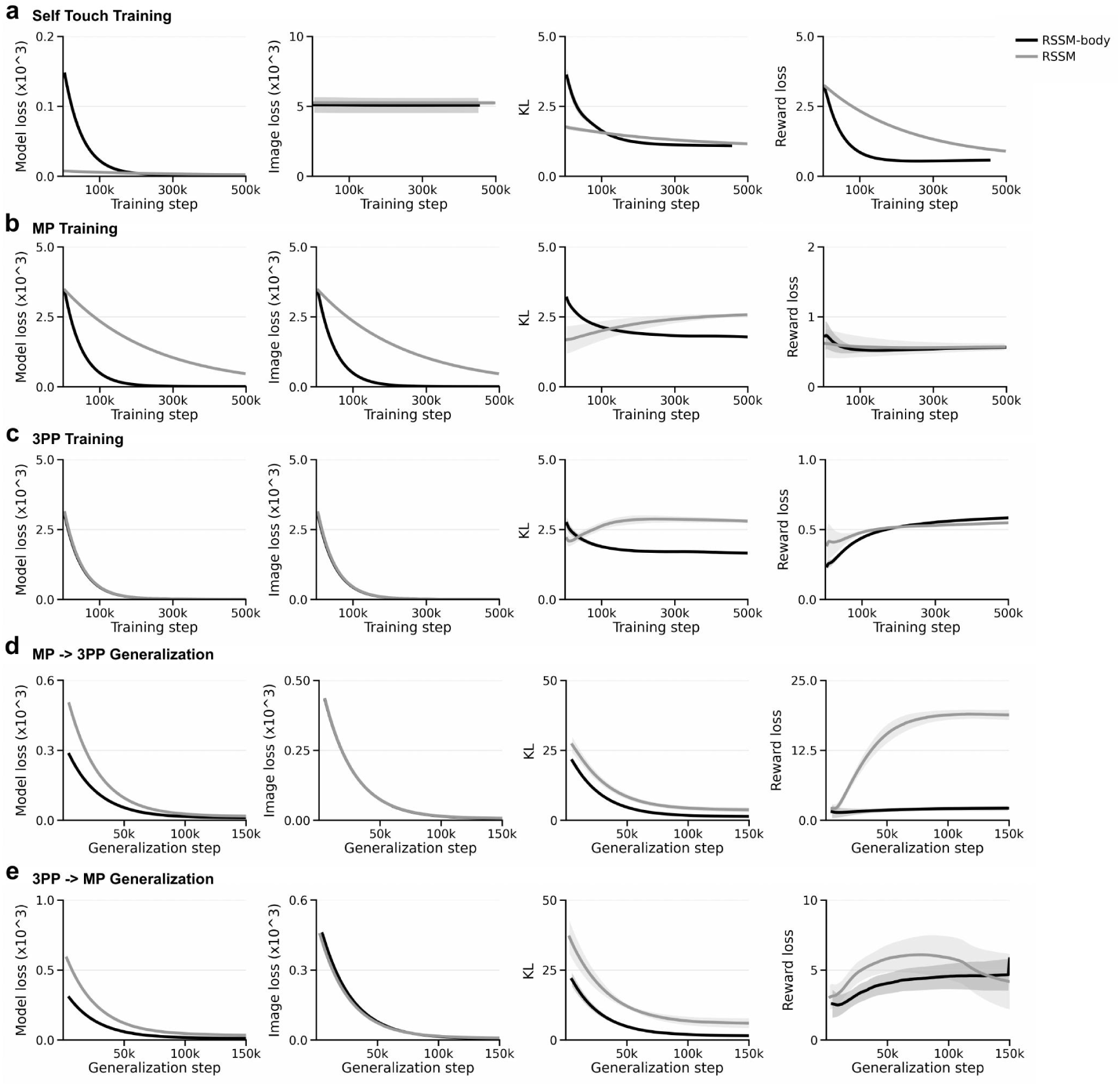
World-model losses during self-recognition training and generalization in robots. **a,** World-model, image reconstruction, Kullback–Leibler (KL) and reward losses during self-touch pretraining without vision. **b,** The same losses during MP target-touch training. **c,** Losses during third-person-perspective (3PP) target-touch training. **d,** Losses during MP→3PP adaptation. **e,** Losses during 3PP→MP adaptation. Black and grey curves show RSSM-body and RSSM, respectively. The auxiliary prediction losses are shown in Fig. 4e,f,k,l and Extended Data Fig. 1e,f.

**Extended Data Fig. 5:**
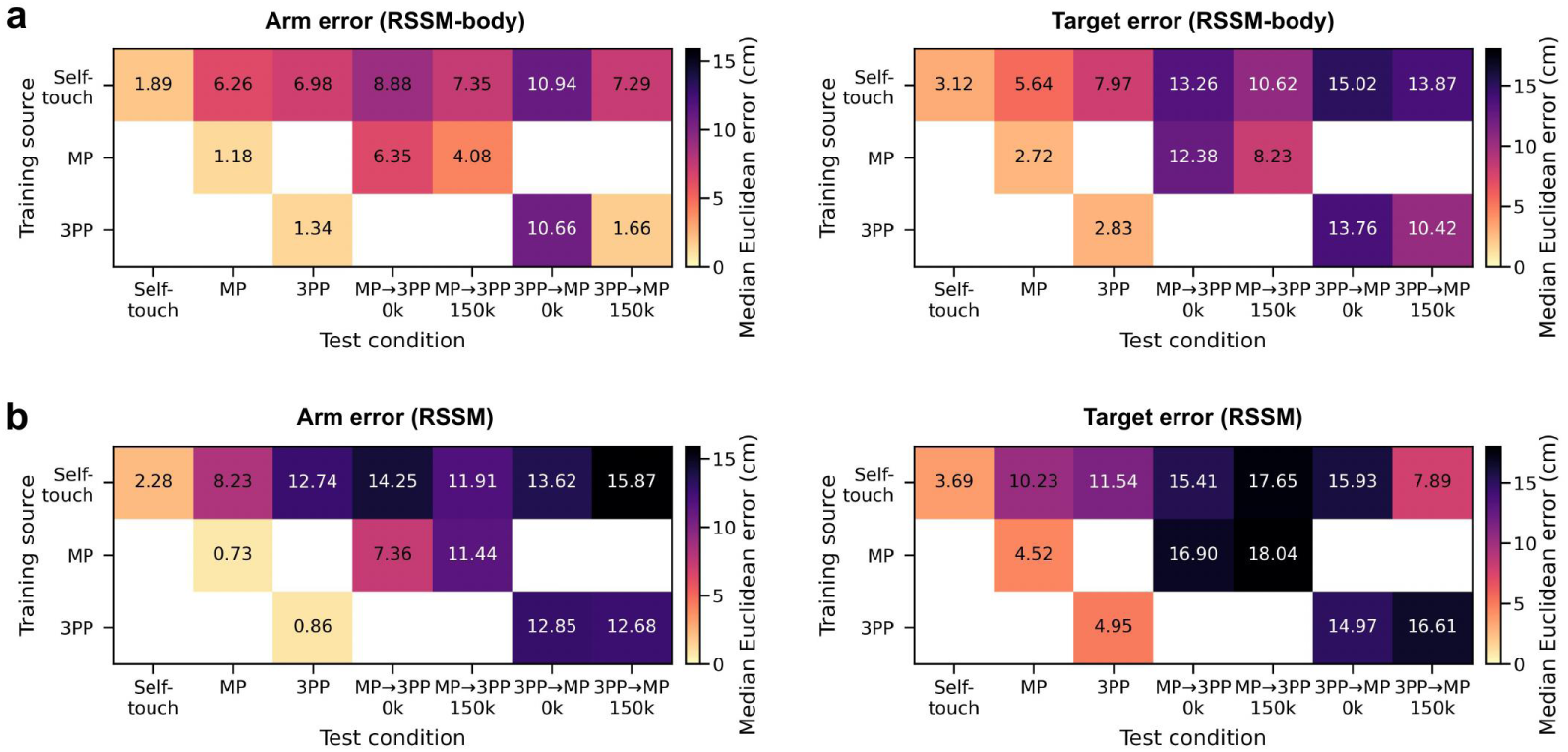
Arm and target decoding errors across stages and scenarios in robots. a,b, Errors of the linear ridge decoders for RSSM-body (a) and RSSM (b), with arm landmarks on the left and targets on the right. Rows show decoder training stages: self-touch, MP and 3PP. Columns show tests in those stages and after each view switch, at 0k and 150k adaptation steps. Decoders were held fixed during testing. For each trial, Euclidean errors were summarised by their median in centimetres. These medians were averaged within each seed and then across seeds. Darker colours indicate larger errors; blank cells indicate combinations not analysed.

**Extended Data Fig. 6:**
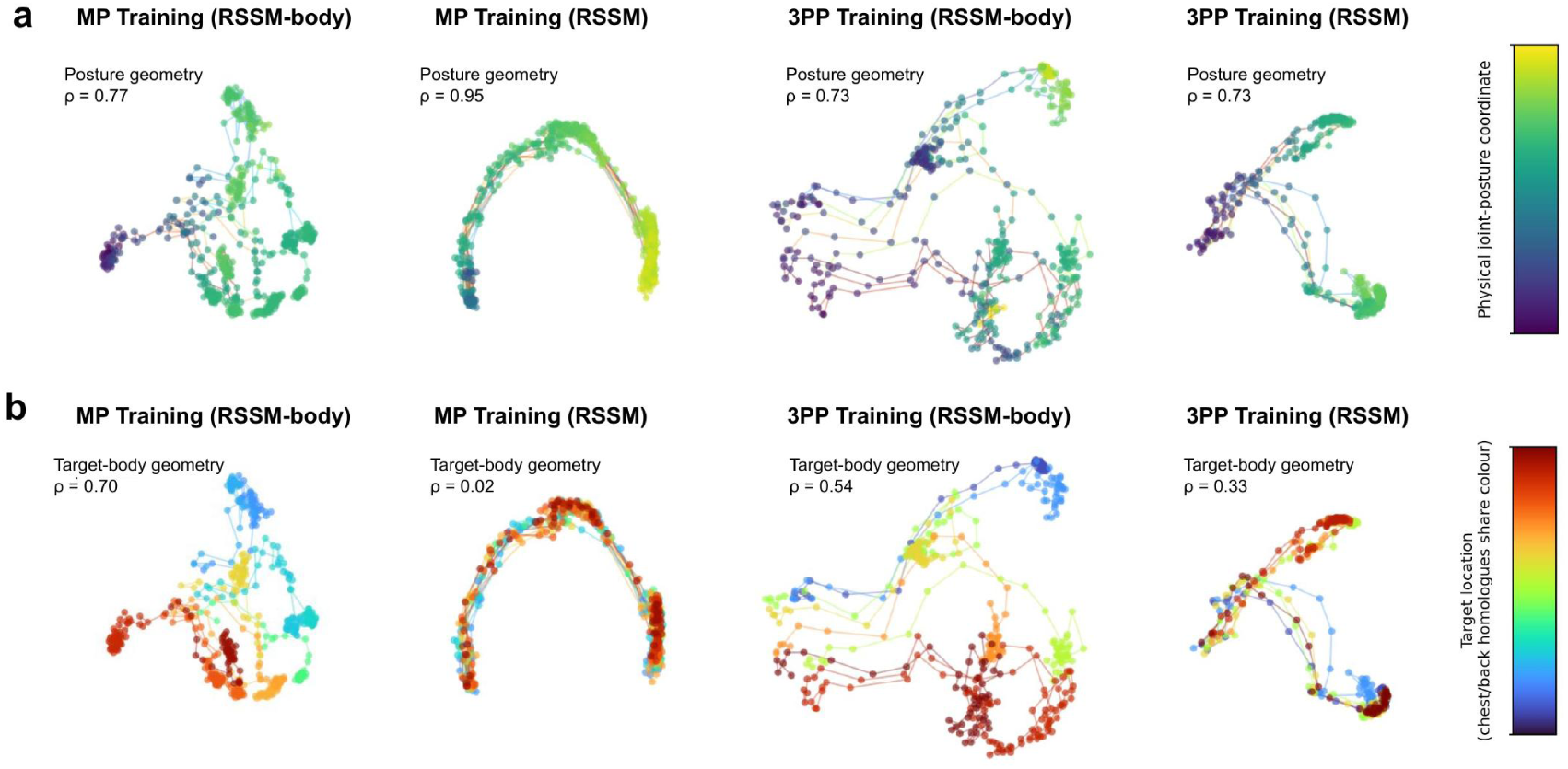
Latent geometry of joint posture and target location during self-recognition training in robots. Columns show RSSM-body and RSSM after MP training (left pair) and 3PP training (right pair), all for seed 0. Two-dimensional multidimensional scaling uses pairwise latent cosine distances. Each point is a sampled frame and lines connect frames from the same trial in temporal order. **a,** Colours show the first principal component of seven joint angles after standardising the angles within each view. Spearman rank correlations (ρ) with distances between standardised postures were 0.77, 0.95, 0.73 and 0.73, from left to right. **b,** The same embeddings coloured by target location, with matching colours for corresponding chest and back grid positions. Spearman correlations with physical distances between targets were 0.70, 0.02, 0.54 and 0.33, from left to right.

**Extended Data Table 1:** Developmental assessment of human infants.

|  |  | **chronological age (months)** | **GDS-C_AQ** | **ASQ_gross motion** |
|---|---|---|---|---|
| all (N=25) | mean | 18.18 | 108.43 | 52.08 |
|  | SD | 3.94 | 19.40 | 8.96 |
| female (N=12) | mean | 18.00 | 103.65 | 52.73 |
|  | SD | 4.08 | 16.29 | 6.84 |
| male (N=13) | mean | 18.35 | 112.84 | 51.54 |
|  | SD | 3.97 | 21.57 | 10.68 |

GDS-C; Griffiths Development Scales-Chinese Edition, AQ; developmental quotient for the Locomotor domain; ASQ: Ages and Stages Questionnaire-3rd Edition.

**Supplementary Video 1 | A macaque transfers mirror self-recognition to a digital mirror view.** (1) Monkey passed MSR test after visual-proprioceptive integration training. (2) Monkey with MSR ability immediately passed light spot and dye mark test in MP scenario. (3) Monkey with MSR ability exhibited dodging and slapping behaviors when seeing virtual spider in MP scenario.

**Supplementary Video 2 | A macaque adjusts its responses after switching to a third-person view.** (1) When switching to 3PP scenario, monkey with MSR ability initially touched the forehead, but adjusted the touching on the back side of its head within 10 seconds (Fig. 1f) when light spot was projected to back side of its head for the first time. (2) When light spot was projected to the waist area of its back for the first time (trial 1), the monkey was unable to touch the light spot (Fig. 1f,g) and displayed apparently exploratory hand movement for reaching the light spot on its back, and could finally touch the spot after several minutes’ exploration.

**Supplementary Video 3 | A macaque passes a third-person mark test after exploration.** In 3PP scenario, monkey with MSR ability spontaneously passed mark test after a period of exploration.

**Supplementary Video 4 | A macaque passes a first-person mark test after the perspective switch.** When switching to 1PP scenario, monkey with MSR ability immediately passed mark test.

**Supplementary Video 5 | A naive macaque learns to respond to marks in a first-person view.** (1) In 1PP scenario, naive monkey could not pass the light spot test on its hands during the first week, but the success rate of grabbing food based on the video image gradually increased (Fig. 2a). (2) By the second week, the naive monkey began to spontaneously recognize that the hands on the video screen were its own, which was confirmed by its success in 1PP mark test (Fig. 2a).

**Supplementary Video 6 | A naive macaque retrieves food shown in a third-person view.** (1) When switching to 3PP scenario, naive monkey failed to find light spot on its back and didn’t exhibit any exploratory behavior. (2) By the second week, the naive monkey was able to obtain the food placed behind it by viewing the video image, but could still not pass mark test in 3PP scenario.

**Supplementary Video 7 | Visual–tactile synchronization training precedes successful third-person mark touching.** (1) Visual-tactile synchronization training in 3PP scenario. (2) Naive monkey passed mark test in 3PP scenario after a few weeks’ training.

**Supplementary Video 8 | A macaque responds to face marks after third-person training.** (1) When switching to MP scenario, monkey with 3PP self-recognition ability immediately passed mark test. (2) Monkey with 3PP self-recognition ability passed MSR test in front of mirror.

## References

1. Botvinick, M. & Cohen, J. Rubber hands ‘feel’ touch that eyes see. Nature 391, 756 (1998).

2. Ehrsson, H. H. The experimental induction of out-of-body experiences. Science 317, 1048 (2007).

3. Lenggenhager, B., Tadi, T., Metzinger, T. & Blanke, O. Video ergo sum: manipulating bodily self-consciousness. Science 317, 1096–1099 (2007).

4. Blanke, O., Slater, M. & Serino, A. Behavioral, neural, and computational principles of bodily self-consciousness. Neuron 88, 145–166 (2015).

5. Amsterdam, B. Mirror self-image reactions before age two. Dev. Psychobiol. 5, 297–305 (1972).

6. Graziano, M. S., Cooke, D. F. & Taylor, C. S. Coding the location of the arm by sight. Science 290, 1782–1786 (2000).

7. Wada, M., Takano, K., Ora, H., Ide, M. & Kansaku, K. The rubber tail illusion as evidence of body ownership in mice. J. Neurosci. 36, 11133–11137 (2016).

8. Fang, W., et al. Statistical inference of body representation in the macaque brain. Proc. Natl Acad. Sci. USA 116, 20151–20157 (2019).

9. Hayatou, Z. et al. Embodiment of an artificial limb in mice. PLoS Biol. 23, e3003186 (2025).

10. Gallup, G. G. Chimpanzees: self-recognition. Science 167, 86–87 (1970).

11. Suarez, S. D. & Gallup, G. G. Self-recognition in chimpanzees and orangutans, but not gorillas. J. Hum. Evol. 10, 175–188 (1981).

12. Gallup, G. G., Jr. Self-awareness and the emergence of mind in primates. Am. J. Primatol. 2, 237–248 (1982).

13. Suddendorf, T. & Butler, D. L. The nature of visual self-recognition. Trends Cogn. Sci. 17, 121–127 (2013).

14. Plotnik, J. M., de Waal, F. B. & Reiss, D. Self-recognition in an Asian elephant. Proc. Natl Acad. Sci. USA 103, 17053–17057 (2006).

15. Reiss, D. & Marino, L. Mirror self-recognition in the bottlenose dolphin: a case of cognitive convergence. Proc. Natl Acad. Sci. USA 98, 5937–5942 (2001).

16. Epstein, R., Lanza, R. P. & Skinner, B. F. “Self-awareness” in the pigeon. Science 212, 695–696 (1981).

17. Prior, H., Schwarz, A. & Güntürkün, O. Mirror-induced behavior in the magpie (Pica pica): evidence of self-recognition. PLoS Biol. 6, e202 (2008).

18. Uchino, E. & Watanabe, S. Self-recognition in pigeons revisited. J. Exp. Anal. Behav. 102, 327–334 (2014).

19. Yokose, J., Marks, W. D. & Kitamura, T. Visuotactile integration facilitates mirror-induced self-directed behavior through activation of hippocampal neuronal ensembles in mice. Neuron 112, 306–318.e8 (2024).

20. Kohda, M. et al. If a fish can pass the mark test, what are the implications for consciousness and self-awareness testing in animals? PLoS Biol. 17, e3000021 (2019).

21. Kohda, M. et al. Further evidence for the capacity of mirror self-recognition in cleaner fish and the significance of ecologically relevant marks. PLoS Biol. 20, e3001529 (2022).

22. Kohda, M., et al. Cleaner fish recognize self in a mirror via self-face recognition like humans. Proc. Natl Acad. Sci. USA 120, e2208420120 (2023).

23. Anderson, J. R. & Gallup, G. G., Jr. Do rhesus monkeys recognize themselves in mirrors? Am. J. Primatol. 73, 603–606 (2011).

24. Anderson, J. R. & Gallup, G. G., Jr. Which primates recognize themselves in mirrors? PLoS Biol. 9, e1001024 (2011).

25. Chang, L., Fang, Q., Zhang, S., Poo, M. M. & Gong, N. Mirror-induced self-directed behaviors in rhesus monkeys after visual-somatosensory training. Curr. Biol. 25, 212–217 (2015).

26. Chang, L., Zhang, S., Poo, M. M. & Gong, N. Spontaneous expression of mirror self-recognition in monkeys after learning precise visual-proprioceptive association for mirror images. Proc. Natl Acad. Sci. USA 114, 3258–3263 (2017).

27. Anderson, J. R. & Gallup, G. G., Jr. Mirror self-recognition: a review and critique of attempts to promote and engineer self-recognition in primates. Primates 56, 317–326 (2015).

28. Toda, K. & Platt, M. L. Animal cognition: monkeys pass the mirror test. Curr. Biol. 25, R64–R66 (2015).

29. Huttunen, A. W., Adams, G. K. & Platt, M. L. Can self-awareness be taught? Monkeys pass the mirror test—again. Proc. Natl Acad. Sci. USA 114, 3281–3283 (2017).

30. Gallup, G. G., Jr. & Anderson, J. R. Self-recognition in animals: where do we stand 50 years later? Lessons from cleaner wrasse and other species. Psychol. Conscious. 7, 46–58 (2020).

31. Prescott, T. J., Vogeley, K. & Wykowska, A. Understanding the sense of self through robotics. Sci. Robot. 9, eadn2733 (2024).

32. Bongard, J., Zykov, V. & Lipson, H. Resilient machines through continuous self-modeling. Science 314, 1118–1121 (2006).

33. Lanillos, P., Pages, J. & Cheng, G. Robot self/other distinction: active inference meets neural networks learning in a mirror. In ECAI 2020 2410–2416 (IOS Press, 2020).

34. Lanillos, P., Dean-Leon, E. & Cheng, G. Yielding self-perception in robots through sensorimotor contingencies. IEEE Trans. Cogn. Dev. Syst. 9, 100–112 (2017).

35. Jazbec, M. et al. Body-swapping experiment with an android robot: investigation of the relationship between agency and a sense of ownership toward a different body. In 2017 IEEE International Conference on Systems, Man, and Cybernetics (SMC) 1471–1476 (IEEE, 2017).

36. Pitti, A., Pugach, G., Gaussier, P. & Shimada, S. Spatio-temporal tolerance of visuo-tactile illusions in artificial skin by recurrent neural network with spike-timing-dependent plasticity. Sci. Rep. 7, 41056 (2017).

37. Iriki, A., Tanaka, M., Obayashi, S. & Iwamura, Y. Self-images in the video monitor coded by monkey intraparietal neurons. Neurosci. Res. 40, 163–173 (2001).

38. Hafner, D., Pasukonis, J., Ba, J. & Lillicrap, T. Mastering diverse control tasks through world models. Nature 640, 647–653 (2025).

39. Howe, M. L. & Courage, M. L. The emergence and early development of autobiographical memory. Psychol. Rev. 104, 499–523 (1997).

40. Rochat, P. Self-perception and action in infancy. Exp. Brain Res. 123, 102–109 (1998).

41. Call, J. & Tomasello, M. Does the chimpanzee have a theory of mind? 30 years later. Trends Cogn. Sci. 12, 187–192 (2008).

42. Krupenye, C., Kano, F., Hirata, S., Call, J. & Tomasello, M. Great apes anticipate that other individuals will act according to false beliefs. Science 354, 110–114 (2016).

43. Ionta, S. et al. Multisensory mechanisms in temporo-parietal cortex support self-location and first-person perspective. Neuron 70, 363–374 (2011).

44. Blanke, O. Multisensory brain mechanisms of bodily self-consciousness. Nat. Rev. Neurosci. 13, 556–571 (2012).

45. Chinn, L. K., Noonan, C. F., Patton, K. S. & Lockman, J. J. Tactile localization promotes infant self-recognition in the mirror-mark test. Curr. Biol. 34, 1370–1375.e2 (2024).

46. Prescott, T. J. & Wilson, S. P. Understanding brain functional architecture through robotics. Sci. Robot. 8, eadg6014 (2023).

47. Tsakiris, M. My body in the brain: a neurocognitive model of body-ownership. Neuropsychologia 48, 703–712 (2010).

48. Prescott, T. J. Robots are not just tools. Connect. Sci. 29, 142–149 (2017).

49. Bryson, J. J. Patiency is not a virtue: the design of intelligent systems and systems of ethics. Ethics Inf. Technol. 20, 15–26 (2018).

